# BAP1 deficient human and mouse uveal melanomas up-regulate a shared EMT pathway

**DOI:** 10.1101/2023.05.24.542173

**Authors:** Roula Farag, Fagun Jain, Anne Nathalie Longakit, Amy Luty, Catherine D. Van Raamsdonk

## Abstract

Monosomy 3 is a negative indicator for uveal melanoma (UM). A key tumor suppressor on chromosome 3 is the deubiquitinase *BAP1*, which usually has a second hit in cases with monosomy 3. Here, we investigated the role of *Bap1* loss in the GNAQ^Q209L^ mouse UM model. We anticipated that the loss of *Bap1* would enhance tumor growth and/or dissemination in a straightforward manner. Instead, our findings revealed a complex relationship between *Bap1* status and tumor behavior. *Bap1^flox/KO^*was associated with a reduction in the number and size of melanocytic lung lesions. *Bap1* heterozygous loss was associated with macroscopic liver lesions. In mice not expressing GNAQ^Q209L^, *Bap1* heterozygous loss caused melanocyte expansion in the choroid, but only in eyes that were structurally abnormal. Finally, transcriptomic analysis identified a shared *BAP1*-associated gene expression program in mouse and human UM that was strongly enriched for epithelial-to-mesenchymal transition (EMT) and apical junction pathways. Together, these findings suggest that Bap1 influences tumor-cell behavior in a manner that depends on gene dosage, tissue context and the surrounding microenvironment.

## INTRODUCTION

Uveal melanoma (UM) is a molecularly distinct subtype of melanoma that arises in the eye, in the choroid, iris or ciliary body [1]. Radiation therapy or enucleation are effective at treating the primary tumor, but up to half of patients will develop metastatic disease, which most often appears in the liver [2]. Patients with distant metastatic disease have a five-year relative survival rate of around 35% [2]. The identification of important recurrent mutations and overactive signaling pathways in this cancer have been identified, which opens up opportunities for targeted therapies. Almost all UMs activate Gα_q/11_ pathway signaling through specific hotspot mutations in the heterotrimeric G protein alpha subunits, *GNAQ* or *GNA11*, or rarely, through activating *PLCB4* or *CYSLTR2* mutations [3,4,5,6]. Gα_q/11_ pathway activation drives cell proliferation and stimulates the mitogen activated protein (’MAP’) kinase pathway, as well as nuclear localization of YAP1 via a Trio-Rho/Rac signaling circuit [3,7,8,9].

Alongside Gα_q/11_ activation, UMs can carry mostly mutually exclusive mutations in *EIF1AX*, *SF3B1* or *BAP1*, the latter of which resides on Chromosome 3 at 3p21.1 [1,10]. It has long been recognized that monosomy 3 is a strong prognostic indicator for UM, posing a significantly increased risk of early metastasis [11]. *BAP1* encodes the BRCA1-associated protein 1 (BAP1) protein, a C-terminal ubiquitin hydrolase functioning as a global epigenetic regulator through its ability to deubiquitinate histone H2A at lysine (K)119. *BAP1* is the most frequently mutated deubiquitinase in human cancer and is a major tumor suppressor [12]. *BAP1* mutations are found in malignant pleural mesothelioma (frequency of 60%), clear cell renal cell carcinoma (15%), intrahepatic cholangiocarcinoma (25%), hepatocellular carcinoma (15%), and thymic epithelial tumors (7%). *BAP1* mutations are observed at a low frequency in breast cancer, colorectal cancer and bladder cancer. In addition, heterozygous copy number loss of the *BAP1* locus was reported in pancreatic carcinoma (25-40%) and promoted metastasis in a *Kras^G12D^* pancreatic cancer mouse model [13].

The effects of *BAP1* loss are complex. Mammalian BAP1 is thought to act mainly as a transcriptional co-activator through histone H2A-K119 deubiquitination, altering chromatin states [14,15,16,17]. Gene expression profiling has found that the depletion or inactivation of *BAP1* induces the up or down regulation of several thousand genes [18]. BAP1 deubiquitinates and/or stably associates with proteins in a number of transcriptional complexes, including HCF1, YY1, OGT, ASXL1/2, and FOXK1/2 [19,20]. BAP1 can also recruit the H3K4 methyltransferase, KMT2C, and the H3K27 demethylase, KDM6A, to gene enhancers, which favors transcriptional activation [21]. In UM, BAP1 may play unique roles compared to other cell types or cancers [14,22].

Our primary goal in this study was to determine whether *Bap1* loss affects melanoma development in mice expressing oncogenic GNAQ^Q209L^ in melanocytes. The *Mitf-cre* transgene is very potent at inducing melanoma in *Cre-loxP* based mouse models [23,24,25]. Therefore we used it to conditionally knock out *Bap1* and drive expression of GNAQ^Q209L^ in this study. It is important to note that *Mitf-cre* mice exhibit microphthalmia, a congenital eye abnormality. The eyes range in severity from poorly formed to nearly normal. Among the latter eyes, overall eye size is reduced by approximately 20%, accompanied by decreased thickness of the neural retina, retinal pigment epithelium (RPE) and choroid, as well as a shortened iris [26]. Variability in eye size is already present at embryonic day (E)13.5, indicating an early developmental disruption [26]. The cause of the microphthalmia is over-expression of *Mitf* in the optic cup from alternative promoters upstream of the melanocyte specific promoter, which drives Cre [26].

We applied a combination of genetic, phenotypic, and transcriptomic analyses, which revealed an unexpected complexity in the relationship between BAP1 and UM progression. Our results suggest a model in which *BAP1* loss alters the state and environmental responsiveness of uveal melanocytes and UM cells, rather than simply increasing proliferation. The consequences of this altered state depend strongly on gene dosage and tissue context. Complete *Bap1* loss does not substantially increase primary tumor growth in the eye in the presence of strong GNAQ activation. However, in a structurally abnormal eye without GNAQ activation, *Bap1* haploinsufficiency promotes melanocyte expansion. In the lung, complete *Bap1* loss is associated with fewer lesions, whereas *Bap1* haploinsufficiency is associated with liver lesions and potentially increased growth of lung lesions in animals that develop liver disease. At the transcriptional level, both mouse and human *BAP1*-deficient UMs show evidence of altered cell adhesion, polarity, and an EMT-like state, which could underlie the phenotypic variability.

## METHODS

### Mouse husbandry and strains

All experiments were carried out under the approval of the Animal Care Committee at the University of British Columbia (animal protocol under Van Raamsdonk C.D., A19-0148). The *Mitf-cre (Tg(Mitf-cre)7114Gsb)* and *Rosa26-floxed stop-GNAQ ^Q209L^ (Gt(ROSA)26Sortm1(GNAQ*)Cvrk)* alleles have been previously described and are maintained on a C3HeB/FeJ genetic background in the lab [25,27,28,29]. The *Bap1* mouse mutant (*Bap1tm1a(EUCOMM)Hmgu/WtsiIeg*) was generated by Helmholtz Zentrum Muenchen GmbH (Hmgu) for the European conditional mouse mutagenesis (EUCOMM) project. It is available at The European Mouse Mutant Archive (EMMA) under ID EM:08891. This ‘knockout-first’ allele design allows for the production of three different alleles, according to the wishes of the researcher. The original “*Bap1^tm1a^*” allele, available at EMMA, contains an IRES:lacZ trapping cassette, neo cassette, and two loxP sites inserted into *Bap1* intron 5 at position 31256950 of chromosome 14 (build GRCm39). A third loxP site is located downstream of Exon 12 at position 31256950. *Bap1^tm1a^* expresses LacZ and disrupts *Bap1* transcription. Removal of the Frt flanked sequences restores *Bap1* expression and creates a Cre-LoxP conditional allele, called the *Bap1^tm1c^* allele, or “*Bap1^flox^*” in this paper. We crossed *Bap1^tm1a^*/+ mice to *PGK-FLP*/+ mice (*Tg(Pgk1-flpo)10Sykr/J*) to generate the *Bap1^flox^* allele. One *Bap1^flox^*/+ mouse was then crossed to a *EIIa-cre*/+ mouse (*Tg(EIIa-cre)C5379Lmgd/J*) to delete exons 6-12 in all cells and create the “*Bap1*^KO^” allele (also known as *Bap1^tm1d^* by EMMA), a germline knockout. The *Bap1^flox^* and *Bap1^KO^* alleles were then separately backcrossed to the C3HeB/FeJ genetic background to segregate each allele from *PGK-Flp* or *EIIa-cre*, respectively, and for all further maintenance. The *Tyr-cre* transgenic line (*Tg(Tyr-cre)1Lru/J*) was purchased from Jackson labs and crossed to either *Rosa26-fs-GNAQ^Q209L^*or *Bap1^flox^/*+.

### Mouse genotyping

Genomic DNA was extracted from ear notches using the DNeasy Blood & Tissue Kit (Qiagen). DNA was amplified using PCR with 50 ng of genomic DNA, 0.25 mM each dNTP, 1 U Hotstar Taq (Qiagen), 1X Hotstar Taq buffer, and 0.5 μM of each primer, in 25 μL total volume. The PCR reaction consisted of 15 minutes at 94°C to activate TAQ, followed by 40 cycles of 94°C (45 seconds), 57°C (45 seconds), and 72°C (1 minute). The *Bap1* related primers used for genotyping are described in **Table S6**. The assays used to genotype for *Mitf-cre* and *Rosa26-floxed stop-GNAQ ^Q209L^* were previously described [23,24].

### Necroscopy and H&E staining

At the experimental endpoint (4, 7 or 20 weeks), mice were euthanized and processed for necroscopy. The skin from the back was removed and the dorsum of the mouse was photographed. The brain was removed, followed by the eyeball(s) when present. Eyeballs were fixed in Davidson’s fixative for 3 hours, rinsed in 1x PBS, followed by 1 hour in 10% formalin, and washed again in 1x PBS, all at 4°C. The liver and lungs were collected and fixed in 10% formalin for 24 hours. Lungs and liver lesions were photographed with a dissecting microscope. Tissue processing and H&E staining was performed by Wax-it Histology Services Incorporated. Sections were taken at 5 μm. In some mice, whole head tissue sections were performed following decalcification.

### Immunohistochemistry (IHC)

Our complete protocol for IHC is published [30]. In brief, 5 μm paraffin sections were first rehydrated into PBS. Antigen retrieval was performed by incubating the slides in 0.6 L of 98°C citrate buffer pH 6.0 (Vector, H-3300) for 10 min, followed by removal from the heat source and cooling for 40 min at room temperature. Next, sections were washed and then placed in a solution of 10% H_2_O_2_ in 1× PBS, which was then heated to 60°C in an oven and held until all pigment was removed (2.5 hours). Sections were washed again, blocked in 5% normal serum in 1x PBS Triton 0.3% for 1 hr at room temperature and then incubated with primary antibodies (anti-CD68 diluted 1:500, Abcam ab125212; anti-S100b diluted 1:1000, Abcam ab5264; anti-RPE65 diluted 1:250, Thermo Fisher Scientific MA1-16578; anti-Ki67 diluted 1:1000, Abcam ab15580; anti-Arg1 diluted 1:50, Cell Signaling Technologies 93668; anti-CD19 diluted 1:1600, Cell Signaling Technologies 90176; anti-CD3 diluted 1:100, Cell Signaling Technologies 78588; anti-Ly6g diluted 1:100, Cell Signaling Technologies 87048) diluted in blocking solution for 1.5 hours at room temperature. Primary antibody was detected using an Elite ABC kit for HRP using DAB with nickel (Vector, SK-4100). DAB color development incubation times were 20 seconds for CD68, S100b, RPE65, Ki67, and Arg1 and 2 minutes for CD19, CD3, and Ly6G.

### Mouse samples used for RNA sequencing

Uveal melanomas were dissected from *GNAQ^Q209L^*/+; *Mitf-cre*/+ mice with normal sized eyes. Two *Bap1* +/+ mice and 3 *Bap1^flox^/+* mutant mice were used to generate 4 samples for each genotype, as follows. For the *Bap1* +/+ samples, uveal melanoma was dissected from four eyes from two mice. 1-WT and 2-WT came from the left and right eyes of the first mouse, while 3-WT and 4-WT came from the left and right eyes of the second mouse. For the *GNAQ^Q209L^*/+; *Mitf-cre*/+; *Bap1^flox^/+* (*Bap1* mutant) samples, uveal melanoma was dissected from four eyes from three mice. 5-Mut and 6-Mut came from the left and right eyes of the first mouse, while 7-Mut and 8-Mut came from the second and third mice dissected. All the mice were female and 20 weeks old.

### Mouse UM RNA sequencing and DE analysis

Following euthanasia, mouse eyes were removed with curved forceps. On ice, each eye was pulled open using forceps and pieces of UM tissue were collected and placed in Trizol (Life Technologies). RNA was isolated according to the manufacturer’s protocol. Quality control of the mouse samples was performed using the Agilent 2100 Bioanalyzer. Samples were then prepped according to the standard protocol for NEBnext Ultra ii Stranded mRNA (New Englad Biolabs). All samples had an RNA Integrity Number (RIN) greater than or equal to 7. The samples were all poly(A) selected. RNA-sequencing was performed on the Illumina NextSeq 500 with Paired End 43bp x 43bp reads. The library consisted of ∼20-25 million reads in total per sample. Illumina’s bcl2fastq2 was used for demultiplexing the sequencing data and these reads were then aligned to the reference genome of *Mus Musculus* mm10 using STAR aligner. The FASTQ files are available at the Sequence Read Archive (SRA) database under project, PRJNA900359 (https://www.ncbi.nlm.nih.gov/sra/PRJNA900359). The aligned read counts were used as input files for Differential Expression (DE) analysis using DESeq2 on R 4.0.3 following the “rnaseqGene” Bioconductor package. We computed log transformations of the read counts to visually identify sample relationships through principal components analysis (PCA) plots and heatmaps. The sample distance heatmap was created using the R function, *dist,* which calculates the Euclidean distance between samples. The rlog and variance stabilizing transformations ensure that all the genes have an equal contribution to the sample distances. We used the raw counts for statistical testing to determine differential expression with DESeq2. The parameters of the DESeq function involve controlling for sequencing depth between samples by estimating size factors and estimating gene dispersion values using negative binomial distribution on the raw gene counts. The DESeq2 package uses Benjamini-Hochberg adjustment to determine the p-adjusted value.

### TCGA human UM dataset analysis

We used htseq count files as input for RNA-seq DE analysis. DE analysis was conducted using DESeq2 on R 4.0.3 following the rnaseqGene Bioconductor package, similar to the RNA-seq analysis for the mouse samples. As with the mouse samples, we filtered the read counts to select genes with read counts that were greater than 10 for 3 or more samples. However, we used variance stabilizing transformation (VST) to stabilize the variance across the range of mean counts. We chose VST for the TCGA data because it works on medium-to-large datasets. VST was used to create the PCA plot and heatmap to visualize the sample-sample distances. Kaplan-Meier plots with univariate Cox proportional hazards were determined using SurvivalGenie [31], an open source online tool available at https://bbisr.shinyapps.winship.emory.edu/SurvivalGenie/

### Statistical test of the overlap in DE genes in mouse and human UM

We first determined the list of genes whose expression was detected in mouse and human UM in the above RNA analyses. We then eliminated all genes that did not have the same name in mouse and human (this does not mean that all remaining genes have the same function in both species). This left 12,258 genes. We then determined whether each gene was differentially expressed in *BAP1/Bap1* mutant versus wildtype UM. DE genes had a FDR < 0.05 with no log_2_FC cutoff. Each gene therefore has one of three outcomes (up-regulated, down-regulated or not DE) in each species. See **Supplementary Table 4** for the designation of every gene in each species. We created a full 3 x 3 table of up/down/not DE genes in human and mouse UM. The Chi-square test was used to test whether mouse and human DE status was independent overall:

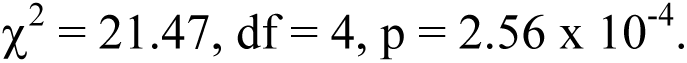

Next, we performed a permutation test. That is, the analysis randomly shuffled the labels (up/down/not DE) for one species 100,000 times and counted the number of concordant genes (DE in the same direction in both species) each time. This analysis determined that the number of concordant genes expected by chance alone is 210.9 +/-13.2 (versus the 270 actually observed). The permutation test count never reached 270 concordant genes. The observed fold enrichment was 1.28x over expected, p = 1 x 10^-5^.

## RESULTS

### *Bap1* mutant alleles in the study

To study the effects of *Bap1* deficiency in mouse UM, we obtained the engineered *Bap1^tm1a^* allele from the European Mouse Mutant Archive (EMMA) [32]. This “knockout-first” allele expresses the LacZ marker from the *Bap1* locus, disrupting *Bap1* transcription (**Figure S1**). Removal of the Frt flanked LacZ sequences restores *Bap1* expression and creates a *Cre-LoxP* conditional allele. We crossed the *Bap1^tm1a^*/+ mice to *PGK-FLPo*/+ mice to generate the conditional *Bap1* floxed allele (“*Bap1^flox^*”). Some *Bap1^flox^*/+ mice were then crossed to *EIIa-cre*/+ mice to delete exons 6-12 in the germline and create a *Bap1* constitutive knockout allele (“*Bap1^KO^*”). *Bap1^flox^*/+ and *Bap1^KO^*/+ lines were each backcrossed to the C3HeB/FeJ genetic background to segregate the alleles from *PGK-FLPo* and *EIIa-cre*, respectively, and were continued to be backcrossed to make the lines inbred. To check that the *Bap1^KO^* allele behaves like a null allele, we intercrossed *Bap1^KO^*/+ mice. *Bap1* constitutive homozygous loss was previously shown to be lethal before the 8th embryonic day [33]. Consistent with this, we obtained no *Bap1^KO/KO^* progeny when genotyping intercross progeny at weaning age (expected frequency = 25%, p = 0.0175, chi square analysis) (**Figure S2**).

### Primary pigmentary phenotypes in *Bap1*-deficient mice expressing GNAQ^Q209L^

The goal of this project was to determine whether *Bap1* loss modulates tumor growth or progression in the *Mitf-cre*; GNAQ^Q209L^ mouse model. To this end, we generated a mouse cohort consisting of *Mitf-cre/+; R26-fs-GNAQ^Q209L^/+* mice [24] with one of four possible *Bap1* genotypes (+/+, *flox*/+, *KO*/+, and *flox/KO*) and aged the mice to 20 weeks. As mentioned, the *KO* allele is constitutively knocked out in all cells from the start of development, while the *flox* allele is conditional and is knocked out only in melanocytes that express *Mitf-M*.

In this cohort, 80% of the mice had severe microphthalmia due to the *Mitf-cre* transgene, such that an intact eyeball could not be located and dissected during necroscopy of the head. However, severe microphthalmia like this does not prevent melanoma growth in the eye socket in this mouse model. Sections of whole heads demonstrated abnormal growth of pigmented tissue within the rudimentary eyes (**Figure 1A**). Such a degree of eye abnormality made it difficult to compare the size of the primary intraocular melanomas by *Bap1* genotype. Of the 9 mice with at least one complete eyeball (“milder mice”), 4 were *Bap1 +/+*, 3 were *Bap1^KO^/*+, and 2 were *Bap1^flox/KO^* (**Figure 1B**). These eyeballs were sectioned and stained with H&E and all exhibited an abnormal overgrowth of pigmented tissue. *Bap1^KO^*/+ did not significantly alter the size of this tissue compared to *Bap1* +/+, but there were not enough mildly affected mice of the other two *Bap1* mutant genotypes to perform statistical analysis (**Figure 1C,D**). Loss of *Bap1* did not increase primary melanocyte overgrowth in the meninges of the central nervous system (**Figure S3, S4**). *BAP1* mutation are found in human primary meningeal melanocytic tumors of the central nervous system [34,35]. No progression to overt, raised tumors outside of the eyes was observed by 20 weeks of age and no mice enrolled in the study were found dead.

**Figure 1.**
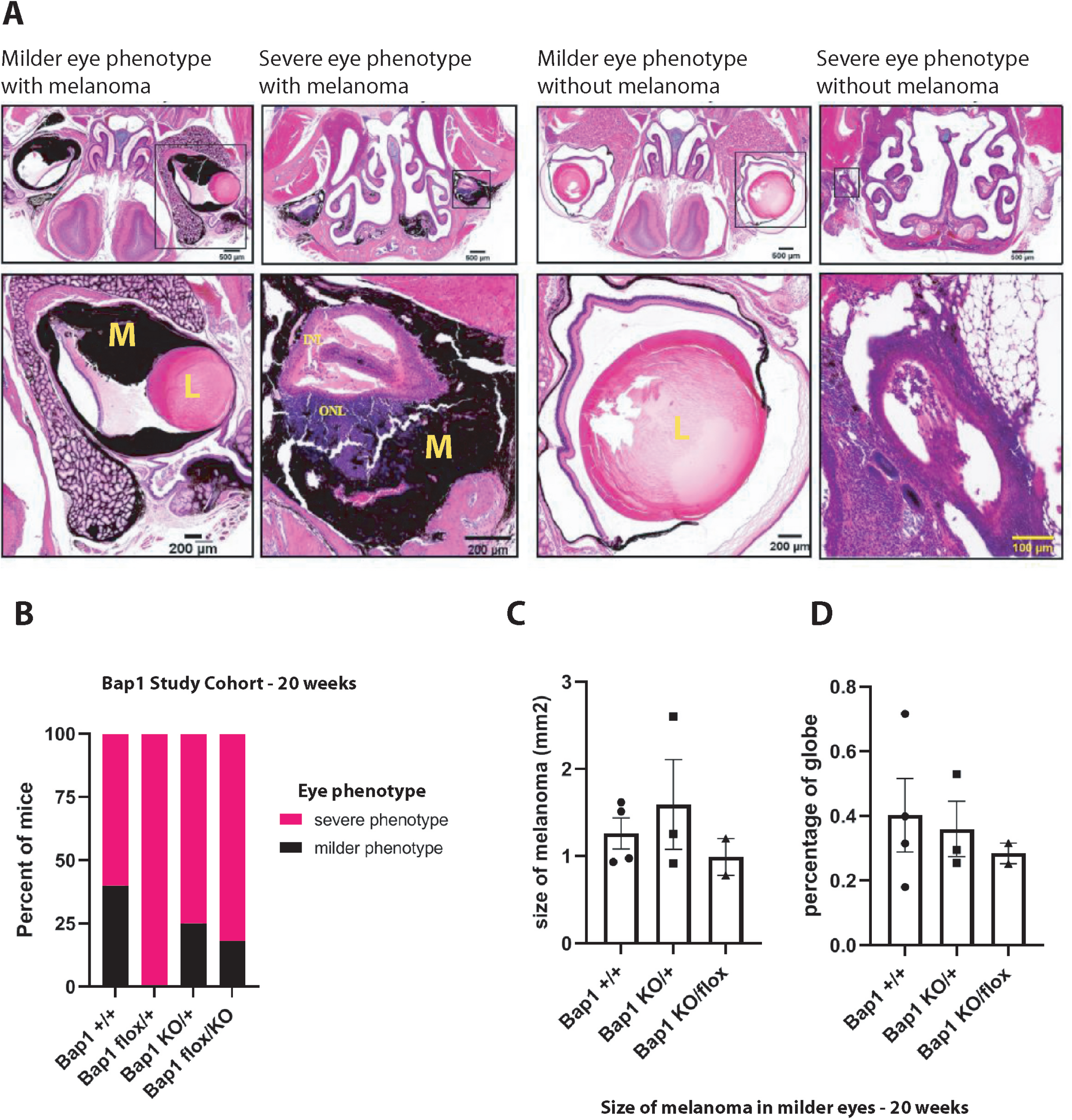
Analysis of eyes in 20 week old *Mitf-cre*; GNAQ^Q209L^ mice with *Bap1* deficiency. **A)** Top row: Representative H&E stained images of sectioned heads of mice with milder or severe microphthalmia, with or without the expression of GNAQ^Q209L^ in melanocytes. Bottom row: Enlargement of an eye in each mouse. Tumor growth was present in all mouse eyes expressing GNAQ^Q209L^. M, melanoma; L, lens; INL, inner nuclear layer of retina; ONL, outer nuclear layer of retina. **B)** Proportions of mice with milder or severe microphthalmia in the 20 week *Mitf-cre*; GNAQ^Q209L^ cohort by *Bap1* genotype. **C)** Quantification of the average area of tumor tissue at the middle of the eye in *Mitf-cre*; GNAQ^Q209L^ mice with milder microphthalmia by *Bap1* genotype. There was no significant difference between the *Bap1^KO^*/+ and +/+ genotypes. **D)** Quantification of the average proportion of the eyeball globe occupied by tumor tissue. There was no significant difference between the *Bap1^KO^*/+ and +/+ genotypes. In C and D, each dot represents the measurement from one mouse. Error bars represent S.E.M.

### Lung lesion analysis in *Bap1* deficient mice expressing GNAQ^Q209L^

The *Mitf-cre*; GNAQ^Q209L^ mouse model exhibits numerous pigmented lung lesions in a broad range of sizes from clusters of a few cells to macroscopic nodules [24] (**Figure 2A**). The origin of these lesions (*e.g.* unknown resident melanocyte population in the lungs, metastases from the UM in the eyes or from another organ) has not been determined. Nevertheless, we were interested in whether *Bap1* loss influences the number or size of the lung lesions. We photographed and measured the area of each lung lesion in whole mount lung preparations from the 20 week cohort (n = 44 mice; 928 lesions total) in a genotype-blinded manner. Average counts per mouse and average size per mouse are plotted in **Figures 2B** and **2C**, respectively.

**Figure 2.**
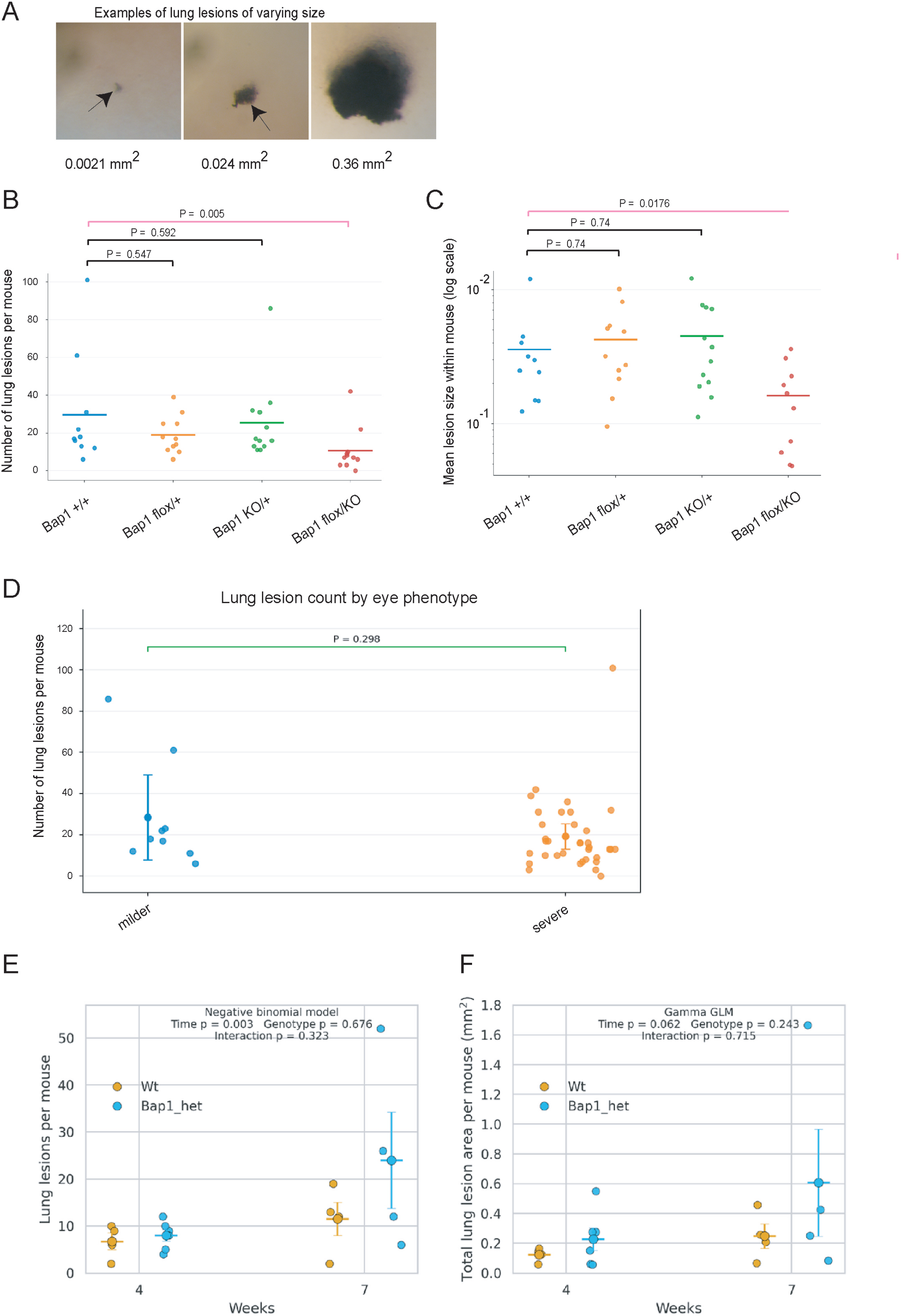
Analysis of lung lesions in the *Mitf-cre*; GNAQ^Q209L^ model with *Bap1* deficiency. **A)** Examples of pigmented lung lesions of increasing size (area indicated in mm^2^) observed in the *Mitf-cre*; GNAQ^Q209L^ model. **B)** Graph of mean lung lesion counts by *Bap1* genotype in 20 week old *Mitf-cre*; GNAQ^Q209L^ mice. Points show individual mice. **C)** Graph of mean lung lesion size by *Bap1* genotype in 20 week old *Mitf-cre*; GNAQ^Q209L^ mice. Points show individual mice. Lesion size is plotted as log scale (mm^2^). **D)** Graph of lung lesion counts by microphthalmia phenotype (milder or severe). Points show individual mice. Error bars show mean +/-95% confidence interval. P values in **B** and **C** compare each *Bap1* mutant genotype to *Bap1* wildtype using negative binomial regression adjusted for eye phenotype. P value in **D** is from negative binomial regression for severe versus milder eye phenotype, adjusted for *Bap1* genotype. **E)** Number of lung lesions per mouse at 4 and 7 weeks in +/+ and *Bap1^flox^***/+** mice. Each point represents one mouse; horizontal lines and error bars indicate mean ± SEM. Lesion number was analyzed using a negative binomial regression model including terms for time (4 vs 7 weeks), genotype, and the time × genotype interaction. **F)** Total lung lesion area per mouse (mm²), calculated as the sum of the areas of all lesions within each mouse. Each point represents one mouse; horizontal lines and error bars indicate mean ± SEM. Total lesion area was analyzed using a Gamma generalized linear model with log link including terms for time, genotype, and the time × genotype interaction.

In a negative-binomial regression model adjusted for eye phenotype, *Bap1^flox/KO^* mice had fewer lung lesions than *Bap1 +/+* mice (adjusted count ratio, 0.37; 95% CI, 0.20–0.69; Holm-adjusted *P* = 0.0049), corresponding to an estimated 63% reduction in lesion number. Neither *Bap1^flox^/+* nor *Bap1^KO^/+* mice differed significantly from *Bap1 +/+* mice in lung lesion count (Holm-adjusted *P* = 0.547 and 0.592, respectively). Mean lesion size was calculated for each lesion-bearing mouse and analyzed using gamma regression adjusted for eye phenotype. Lesions in *Bap1^flox/KO^* mice were significantly smaller than those in *Bap1 +/+* mice (adjusted size ratio, 0.46; 95% CI, 0.27–0.80; Holm-adjusted *P* = 0.0176), representing an estimated 54% reduction in mean lesion size. No significant differences in lesion size were observed for either heterozygous *Bap1* genotype relative to *Bap1 +/+*. Eye phenotype was not independently associated with lung-lesion number: mice with severe microphthalmia had an adjusted lesion-count ratio of 0.75 relative to mice with milder microphthalmia (95% CI, 0.43–1.29; *P* = 0.298) (**Figure 2D**).Together, these findings indicate that *Bap1^flox/KO^* is unexpectedly associated with a reduced lung lesion burden in this model.

### The number of lung lesions increases between 4 and 7 weeks in the *Mitf-cre*; GNAQ^Q209L^ mouse model

We next bred a second cohort of *Mitf-cre;* GNAQ^Q209L^ mice with either *Bap1 +/+* or *Bap1^flox^/+* genotypes and only included mice born with mild eye phenotypes. We counted and measured the lung lesions in mice euthanized at either 4 weeks or 7 weeks old. The number of lung lesions per mouse was significantly higher at 7 weeks than at 4 weeks (negative binomial regression, p = 0.003) (**Figure 2E**). The main effect of *Bap1* genotype was not significant (p = 0.676), and there was no significant time × genotype interaction (p = 0.323), indicating that both genotypes showed a similar increase in lesion number over time. Total lung lesion area per mouse, calculated by summing the area of all lesions within each mouse, also increased from 4 to 7 weeks, but with suggestive statistical significance (Gamma generalized linear model with log link, p = 0.062) (**Figure 2F**). These data show that lung lesion burden increases between these two time points, primarily through an increase in lesion number.

### Liver lesions in *Bap1* heterozygous mice expressing GNAQ^Q209L^

We carefully checked every liver for lesions in all mice while blinded to genotype. In the first cohort, macroscopic liver lesions were found in 1/11 (9%) of the *Bap1^flox^*/+ mice and 2/12 (17%) of the *Bap1^KO^*/+ mice, but not in *Bap1* +/+ (n=10) or *Bap1^flox/KO^* (n=11) mice (**Figure 3A**). In each case, there was only one lesion in the liver. In the second round of breeding in another subset aged to 20 weeks, we found a macroscopic liver lesion in one *Bap1^flox^*^/^+ mouse out of 5, with no liver lesion in 7 *Bap1* +/+ control animals from the same breeder pairs. In addition, no liver lesions had been found in the 19 *Mitf-cre*; GNAQ^Q209L^ mice that we described previously [24]. A contingency table analysis of all *Bap1* heterozygous mice versus *Bap1* +/+ mice found a significant difference in macroscopic liver lesion frequency (p=0.0322, Fisher’s Exact test, **Figure 3B**). No macroscopic lesions were found in the livers of the mice aged to only 4 or 7 weeks of age.

**Figure 3.**
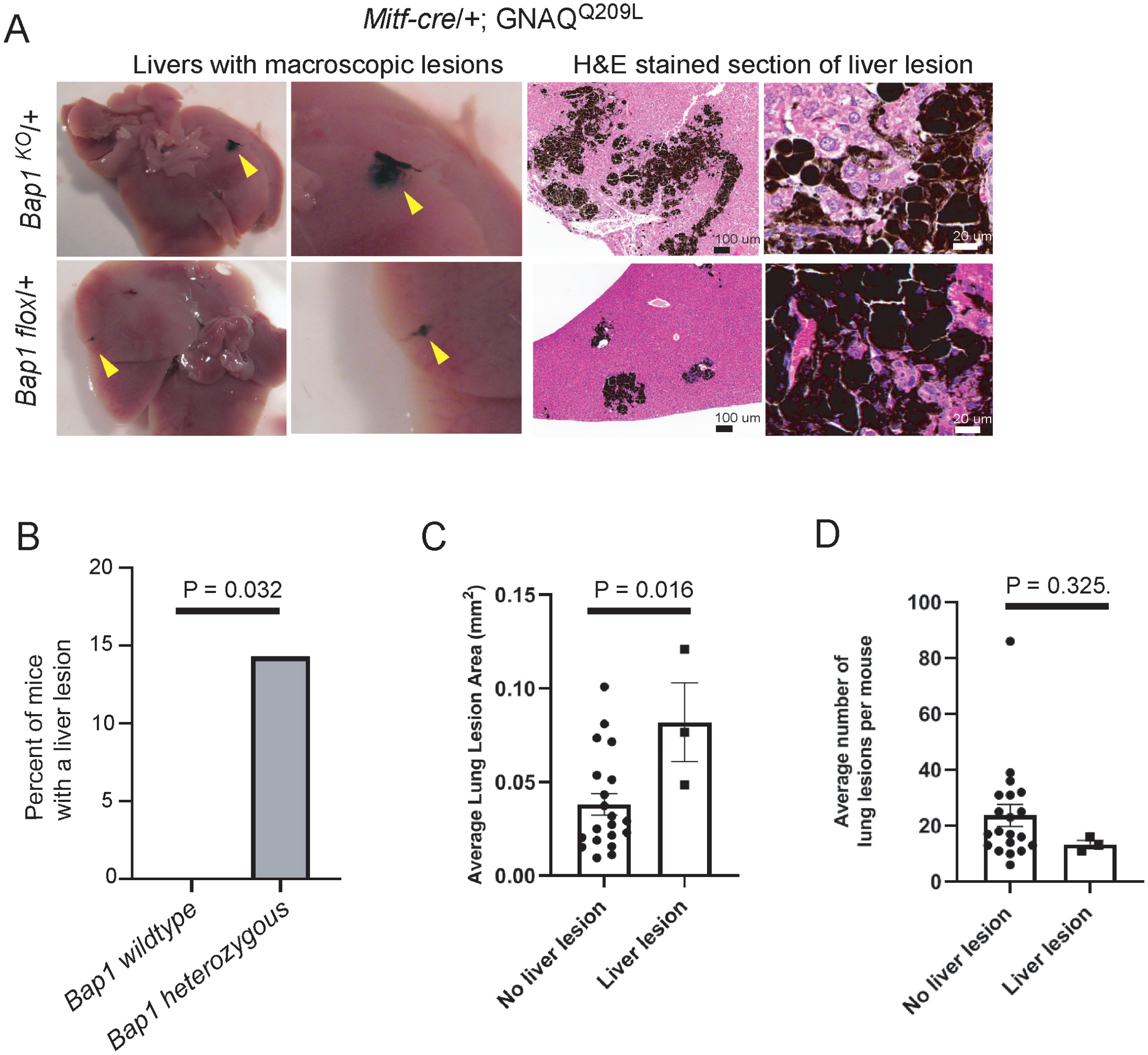
Melanocytic liver lesions in *Bap1* heterozygous *Mitf-cre*; GNAQ^Q209L^ model. **A)** Images on left side show macroscopic liver lesions in *Mitf-cre*/+; *R26-fs-GNAQ^Q209L^*/+ mice that are either *Bap1^KO^*/+ or *Bap1^flox^*/+. On the right hand side are H&E stained sections of these lesions. **B)** Graph of the percentage of mice with a macroscopic liver lesion. n= 36 *Bap1* +/+ mice, n=28 *Bap1* heterozygous mice. Affected mice had only 1 liver lesion each. **C)** Graph showing the size of lung lesions in *Bap1* heterozygous mice with or without a liver lesion. Each point represents the average lung lesion size in one mouse. **D)** Graph showing the number of lung lesions in *Bap1* heterozygous mice with or without a liver lesion. Each point represents the number of lung lesions in one mouse.

Pearson *et al.* demonstrated that liver metastases can actively promote systemic cancer progression by reprogramming the immune environment [36]. To examine whether this could play a role in the *Mitf-cre*; GNAQ^Q209L^ mouse model, we compared the average size of lung lesions in the three mice with liver lesions with the remaining *Bap1* heterozygous mice without a liver lesion at 20 weeks of age. In the mice with liver lesions, there was a significant increase in average lung lesion size per mouse (p=0.016, unpaired ttest, **Figure 3C**), with no difference in the number of lung lesions per mouse (p=0.326, unpaired ttest, **Figure 3D**). All three mice with a liver lesion had a severe eye phenotype.

We performed H&E staining and IHC (using melanin bleaching) [30] to examine the expression of S100B (melanoma), Ki67 (cell proliferation), CD68 (macrophages), Arg1 (myeloid derived suppressor cells and hepatocytes), Ly6G (neutrophils/granulocytes), CD3 (T cells), and CD19 (B cells) in the liver. What appeared macroscopically to be single lesions presented as multiple separate clusters in sections (**Figure 3A**). The lesions exhibited what is called a replacement growth pattern, as single S100b-positive tumor cells could be found intermingled with hepatocytes on the edges of the lesions [37] (**Figure S5A**). Lesions were accompanied by abundant CD68-positive macrophages, which were intensely dark with phagocytosed melanin and had abundant cytoplasm (**Figures 4A,B, Figure S5A,B**). The intensity of CD68 expression varied among macrophages, even between neighboring cells, suggesting heterogeneity in lysosomal activity or maturation state (**Figure 4B**). Some macrophages were organized into unusual looking, tightly packed clusters, often with a circling layer of S100b-positive cells around the periphery (**Figure 4C**). In some of these tightly packed clusters, we observed cells in the interior that were Ki67-positive, suggesting cell activation; however, it’s not possible to tell the identity of the cells with the Ki67-positive nuclei (**Figure 4D**). Using their distinctive morphology to identify them, we found no evidence that the large, round macrophages were Arg1-positive (**Figure 4G**).

**Figure 4.**
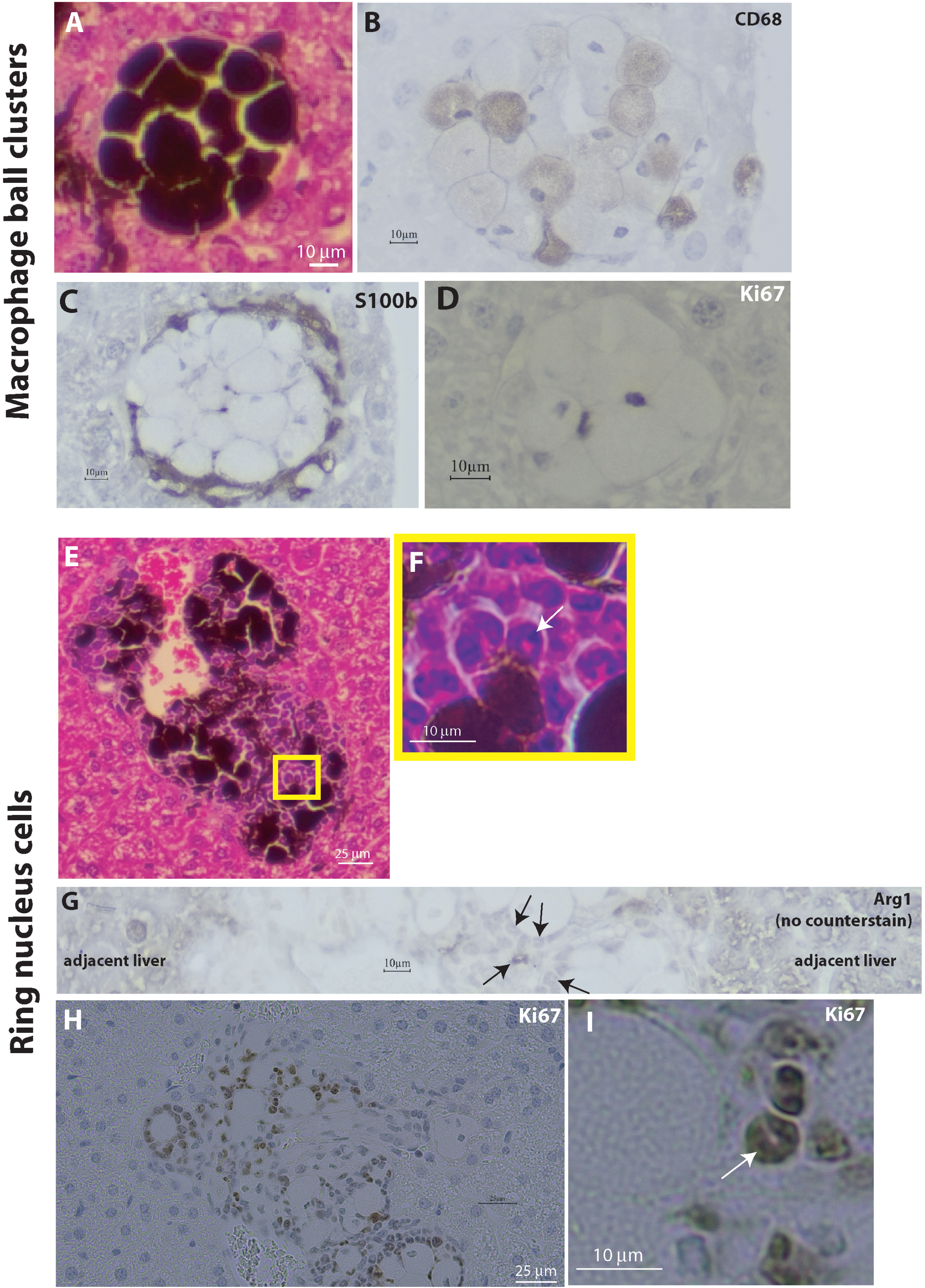
Immune cells in *Bap1* heterozygous *Mitf-cre*; GNAQ^Q209L^ liver lesions. **A)** Ball of macrophages stained with H&E, showing abundant cytoplasm and intense melanin. **B-D)** IHC detection of CD68 (B), S100b (C), and Ki67 (D) in sections of macrophage balls, with melanin bleaching and hematoxylin counterstaining. **E,F)** H&E stained section showing compact hyperchromatic cells with basophilic ring shaped nuclear material. The clearest ring shaped nucleus is indicated with a white arrow. **G**) IHC detection of Arg1 expression in a section of liver containing a lesion, following melanin bleaching, but without hematoxylin counterstaining. Normal hepatocytes express Arg1 on either side of the lesion. Arg1 expression is also detected in a cluster of cells with ring shaped nuclei in the center (black arrows). **H,I**) IHC detection of Ki67 in an area with many cells exhibiting ring shaped nuclei, following melanin bleaching and with hematoxylin counterstaining. A strongly Ki67-positive cell with this morphology is indicated with a white arrow.

In the lesions, we also observed an interesting cell type that was compact/small and hyperchromatic, with a very basophilic ring of nuclear material (**Figure 4E,F**). Cells with this morphology were Arg1-positive (**Figure 4G**) and Ki67-positive (**Figure 4H,I**). We observed no positivity for CD3 in cells with this morphology. Given the Arg1 and Ki67 positivity, one possibility is that these cells might be a proliferating immature myeloid population. For example, myeloid derived suppressor cells (MDSCs) can secrete or display Arg1 on their surface, which has an immune suppressing effect on T cells (reviewed in [38]).

In terms of other immune cell types, CD19-positive cells were not detected in either the normal liver areas or the lesions (**Figure S5C**), but were detected as expected in control mouse spleen (spleen not shown). Ly6G-positive cells were not present in the normal liver areas and were rarely found in the lesion area (**Figure S5D**). CD3-positive cells were present throughout the liver as expected and were not noticeably concentrated in the lesions (**Figure S5E**).

### *Bap1* heterozygous deficiency in structurally abnormal eyes generates melanocyte hypertrophy

As part of our breeding strategies, we also obtained *Bap1* mutant mice not expressing GNAQ^Q209L^. Interestingly, we noticed that *Mitf-cre*/+; *Bap1^flox^/+* eyes displayed a significant expansion of pigmented tissue relative to *Mitf-cre/+*; +/+ controls (**Figure 5A-E**; p = 0.0025, unpaired t-test). In contrast, no pigmentation abnormalities were observed in the *Bap1^KO^*/+ eyes (without *Mitf-cre*) (**Figure 5F,G**). To confirm that it was melanocytes that were affected in the *Mitf-cre*/+; *Bap1^flox^/+* eyes, we performed immunohistochemistry on bleached sections using markers distinguishing uveal melanocytes (S100b) from RPE cells (RPE65) [30]. The expanded pigmented regions were S100b-positive and RPE65-negative (**Figure 5D**), demonstrating that *Bap1^flox^* promotes the expansion of uveal melanocytes in *Mitf-cre* eyes. Furthermore, *Tyr-cre*/+; *Bap1^flox^/+* eyes did not exhibit the same phenotype (**Figure S6**) and *Mitf* was not over-expressed from the *Mitf-cre* transgene in uveal melanocytes (**Figure S7**).

**Figure 5.**
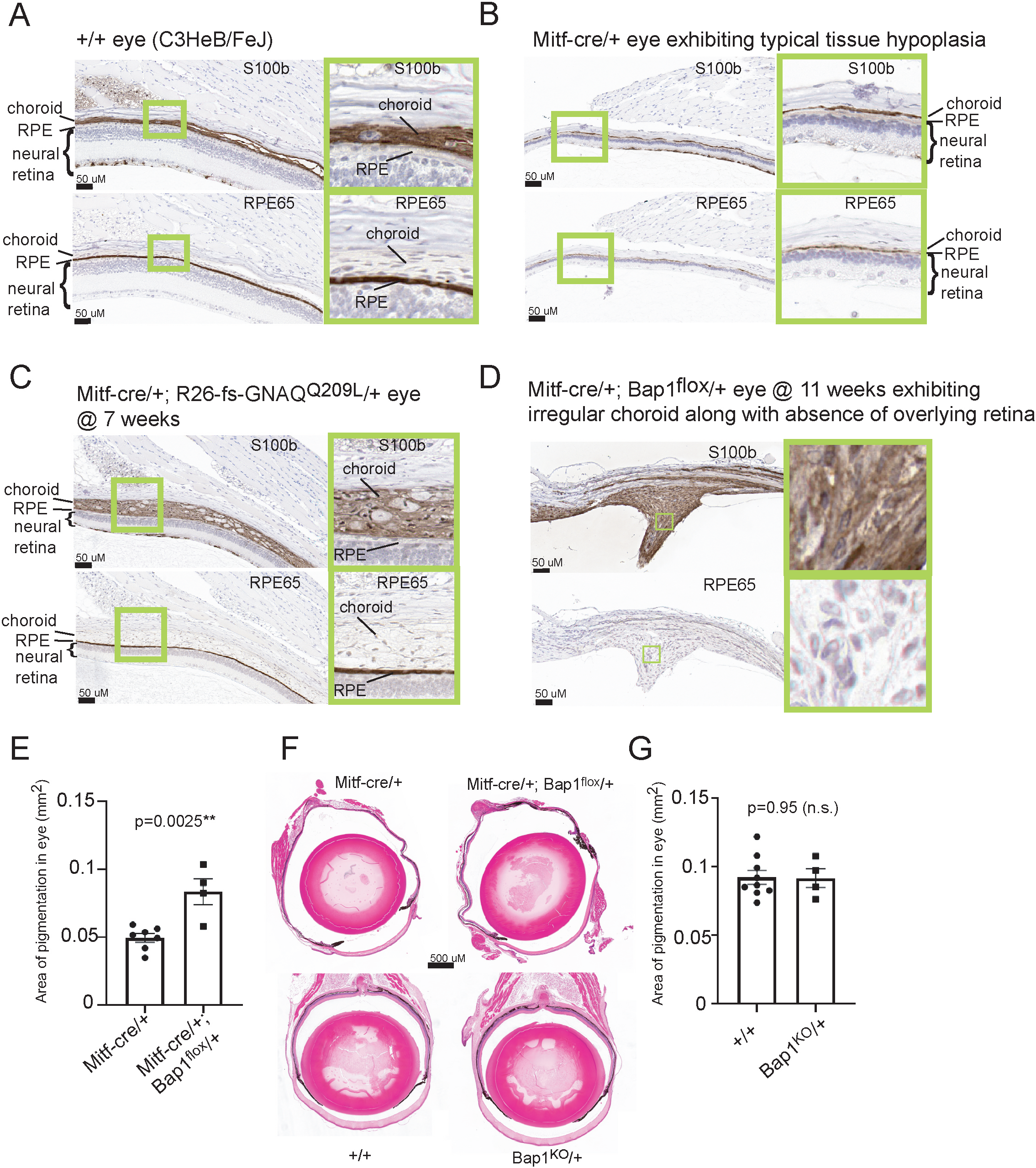
*Bap1* heterozygous deficiency in structurally abnormal eyes generates melanocyte hypertrophy A-D) IHC detection of S100b and RPE65 protein in the eyes of mice of the indicated genotypes. S100b is a marker for melanocytes and melanoma, and is specific for the uveal tract, while RPE65 is specific to the RPE. Sections shown were bleached to remove melanin prior to staining. Green boxes show areas enlarged to the right side of each panel. The abnormal tissue in Mitf-cre/+; Bap1flox/+ eyes is positive for S100b and negative for RPE65 (D). E) Graph shows a significant increase in the area of pigmented tissue in the eyes of Mitf-cre/+; Bap1flox/+ mice compared to Mitf-cre/+; +/+ mice, measured in sections taken at the middle of the eye. Each dot represents the measurement from one eye. Error bars show the S.E.M. F) Representative eyes of the indicated genotypes, unbleached and stained with H&E. G) Graph shows no significant difference in the amount of pigmented tissue in the eyes of Bap1KO/+ mice compared to +/+ mice, measured in area of sections taken at the middle of the eye. Each dot represents the measurement from one eye. Error bars show the S.E.M.

*Mitf-cre* causes thinning of the RPE and neural retina, which likely alters the mechanical and structural constraints imposed on the choroid, where melanocytes reside [26,39]. We hypothesize that this altered microenvironment permits increased proliferative or invasive behavior of *Bap1* heterozygous melanocytes. Consistent with this model, the largest abnormal area we found lacked an overlying retina and Bruch’s membrane (**Figure 5D**). This melanocyte growth pattern may be similar to how human UMs seem to expand more easily in the subretinal space after they breach Bruch’s membrane, acquiring a “mushroom” shape [40]. Taken together, these findings support a model in which loss of *Bap1* enhances the proliferative and/or invasive capacity of uveal melanocytes in the eye, but only in a permissive microenvironment where normal tissue architecture is compromised. This suggests that *Bap1* functions as a context-dependent regulator of normal uveal melanocyte growth.

### RNAseq analysis of *Bap1* haploinsufficiency in mouse UM

We sequenced RNA from UM tissue dissected from mildly affected eyes of *Mitf-cre*; GNAQ^Q209L^ mice that were either *Bap1* wildtype (UM from 4 eyes/2 mice) or *Bap1^flox^/+* (UM from 4 eyes/3 mice) at 20 weeks of age. All mice were female. We examined *Bap1* transcripts using a sashimi plot to visualize splice junctions (**Figure S8**). The only abnormal reads that were present connected exon 5 to exon 13 in the *Bap1^flox^/+* mutant group (10/67 reads), but not in the *Bap1* wildtype group (0/158 reads), as expected from the placement of the *loxP* sites, which flank exons 6-12. To determine how the mouse samples clustered together, we performed exploratory analysis using principal component analysis (PCA) (**Figure 6A**) and a sample distance heatmap (**Figure 6B**). Both analyses showed an unexpected split clustering of the 1-WT and 2-WT samples (left and right eyes of one *Bap1* wildtype mouse) from the 3-WT and 4-WT samples (left and right eyes of a second *Bap1* wildtype mouse). This suggests that there could be higher tumor to tumor variability in the *Bap1* wildtype tumors. DE analysis found that there were 2566 differentially expressed (DE) genes with a p-adj value < 0.05 (**Table S1**). 1232 genes were down-regulated and 1334 genes were up-regulated in the *Bap1^flox^/+* UM samples compared to *Bap1* wildtype UM samples.

**Figure 6.**
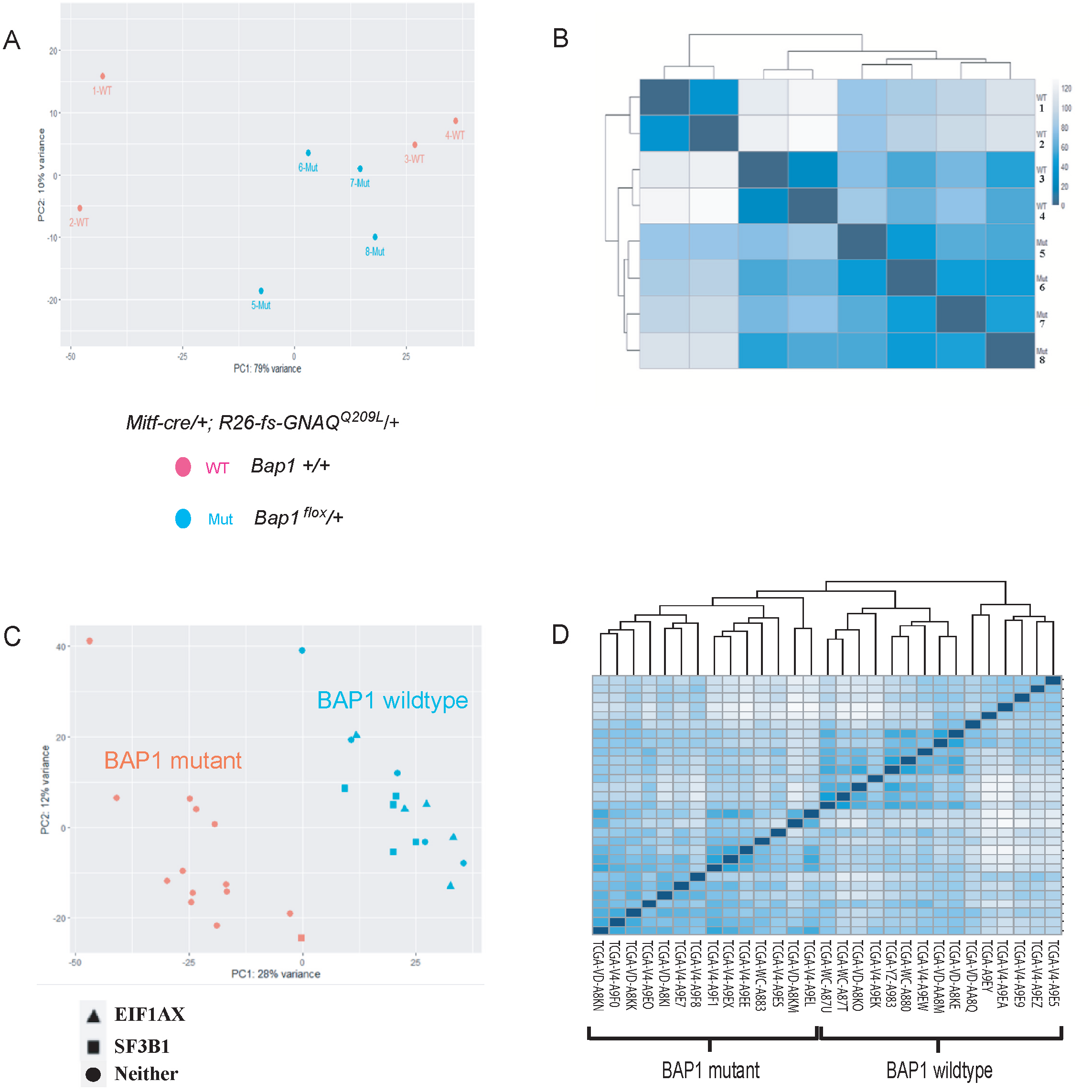
Mouse and human samples used for RNAseq. A,B) PCA plot (A) and sample distance heatmap (B) of *Mitf-cre/+; R26-fs-GNAQ^Q209L^/+*mouse UM samples that were either *Bap1* +/+ (“WT”, pink dots #1-4) or *Bap1^flox^*/+(“Mut”, blue dots #5-8), collected at 20 weeks of age. WT 1 and WT 2, each from a different eye of the same mouse, clustered farther away from the other WT samples than expected. **C,D)** PCA plot (C) and sample distance heatmap (D) of human UM samples from the TCGA-UVM dataset used for DE analysis. Twenty-nine cases in the TCGA-UVM dataset have activating mutations in *GNAQ* and were selected. Fifteen were disomic for chromosome 3 and had no focal mutations in *BAP1* (“*BAP1* wildtype” blue symbols). Of the “*BAP1* mutant” (orange symbol) cases, ten carried monosomy 3 with a *BAP1* alteration detected by either DNA or RNA sequencing. Three samples had monosomy 3 only with no other alteration to *BAP1* reported. One was disomic for chromosome 3 with *BAP1* loss of heterozygosity (LOH) carrying a mutation in *BAP1*. All *BAP1* mutant cases clustered together.

### RNAseq analysis of *BAP1* loss in human UM

We were curious about how the changes in gene expression in the mouse UM model correlated with human UM. We obtained the publicly available UM patient data (TCGA-UVM) [1] from The Cancer Genome Atlas (TCGA). To match the mouse model as closely as possible, we selected the 29 UM patient samples with activating *GNAQ* mutations for study (**Table S2**). We didn’t include the *GNA11* mutant samples because *GNA11* may not act exactly like *GNAQ* in UM: *GNA11* mutations are enriched in human UM metastases compared to *GNAQ* mutations, only *GNAQ* mutations are abundant in benign blue nevi, and *GNA11* mutations are associated with increased risk of death [3,4,41].

The mutational analysis of the TCGA UM dataset previously done by Robertson *et al* [1] using whole exome DNA sequencing and RNA sequencing are repeated here for reference in **Table S2**. Fifteen of the tumors were disomic for chromosome 3 and had no focal mutations in *BAP1*. These were designated as *BAP1* wildtype. Five of these had an *EIF1AX* mutation, 5 had a *SF3B1* mutation and 5 had neither of these genes mutant. The following 14 samples were considered *BAP1* mutant: 10 samples with monosomy 3 and a focal mutation in *BAP1*, one sample with isodisomy 3 carrying a focal mutation in *BAP1,* and 3 samples with monosomy 3, but no focal mutation in the *BAP1* gene. One of the tumors with a *BAP1* focal mutation also had a *SF3B1* mutation, which was the recurrent *R625H* missense.

Using RNAseq data, both the PCA plot and sample distance heatmap that we generated showed a clear separation between the *BAP1* wildtype and *BAP1* mutant UM samples (**Figure 6C, 6D**). We performed DE analysis using methods consistent with the analysis in mice. There were 3056 differentially expressed genes with a p-adj value < 0.05. 1328 genes were down-regulated and 1728 genes were up-regulated in the *BAP1* mutant samples (**Table S3**). Hence, in both mouse and human UM, there was a very large number of differentially expressed genes and slightly more up-regulated than down-regulated genes in *BAP1* mutants.

### Shared transcriptional changes in *BAP1* mutant UM identify EMT and AJ pathways

The thousands of DE genes that are produced when *BAP1* is mutant in either the mouse or the human makes investigating the effects of *BAP1* loss quite challenging. We thought it might be helpful to identify the overlap between the mouse and human results. 12,258 genes with the same name were expressed in both the human and mouse UM samples. 8200 of these genes (67%) were not DE in either species, 3580 (29%) were DE in one species but not the other, and the remaining 478 genes (4%) were DE in both species (genes are listed in **Table S4**). 270 were differentially expressed in the same direction in both mouse and human. This included 162 up-regulated and 108 down-regulated genes (**Figure 7A**). 270 concordantly differentially expressed genes is significantly greater than expected by chance (permutation test, p < 10^-5^), with a 1.28-fold enrichment over the expected ∼211 genes.

**Figure 7.**
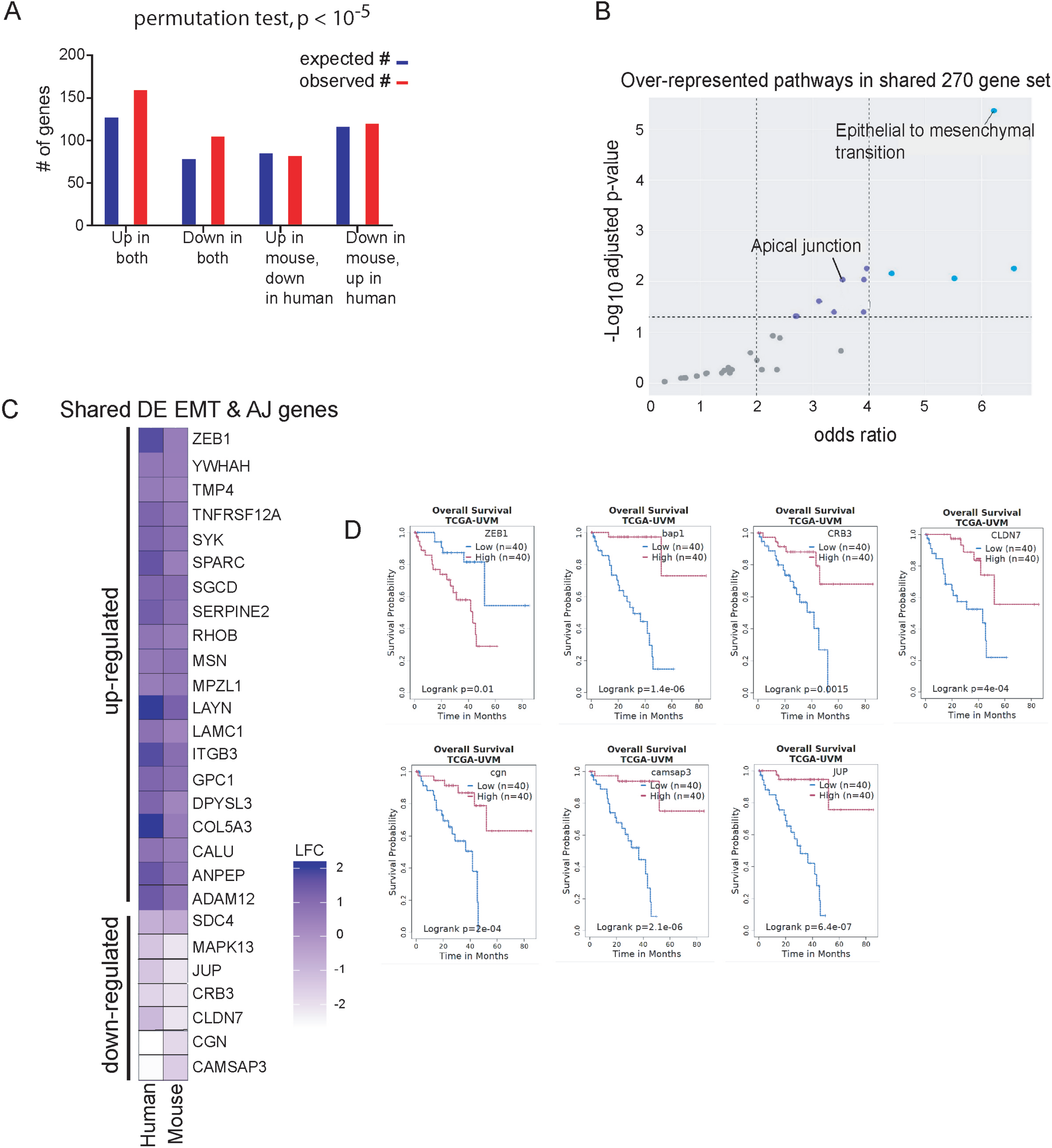
*BAP1* loss is associated with changes to EMT and AJ gene expression in mouse and human UM. **A)** Graph shows the number of genes that are DE in the same or different directions (up or down) in mouse and human UM. Compared to the number expected by random chance, there are significantly more genes that are DE in the same direction in both species (concordant). Counts involving genes that are not DE in one or both species are not shown on this graph. UP in both: 162 observed, 130 expected; DOWN in both: 108 observed, 81 expected; UP in mouse and DOWN in human: 85 observed, 88 expected; DOWN in mouse and UP in human: 123 observed and 119 expected. **B)** Pathway analysis of the 270 shared DE genes in mouse and human, showing that EMT is the most significantly up-regulated pathway. Grey dots are below the significance threshold, while blue dots are the most significant. All pathways, genes and p-values and odds ratios are listed in Supplementary Table 5. **C)** Heatmap showing the log_2_ fold change in shared DE genes in the EMT and AJ hallmarks. **D)** Kaplan-Meier plots showing that low expression of the AJ genes, *JUP*, *CLDN7*, *CRB3*, *CGN* and *CAMSAP3,* or high expression of the EMT inducer *ZEB1,* are significantly associated with reduced survival in the TCGA UVM dataset, along with *BAP1*.

We performed pathway analysis using EnrichR to look for MSigDB Hallmark gene sets _2020 enriched terms in the 270 genes. There were 18 significantly enriched terms (**Table S5**), but by far the most significant was Epithelial to Mesenchymal Transition (EMT) (p-adj = 4.34 x 10^-6^) (**Figure 7B**). EMT is a hallmark of cancer. The shared DE markers of EMT in UM are *ADAM12*, *ANPEP*, *CALU*, *COL5A3*, *DPYSL3*, *GPC1*, *ITGB3*, *LAMC1*, *RHOB*, *SDC4*, *SERPINE2*, *SGCD*, *SPARC*, *TNFRSF12A*, and *TPM4*. All but one of these 15 genes are up-regulated in the *BAP1* mutants, consistent with their known roles promoting EMT (**Figure 7C**). Although it was not a member of the EMT Hallmark gene set, *ZEB1* encodes a major EMT inducer and it was also up-regulated in the *BAP1* mutants.

A second significant term in the analysis that is closely related to EMT is Apical Junction (AJ), supported by 9 genes, none of which is also on the EMT list (p-adj = 0.009) (**Figure 7B,C**). These genes were *CLDN7*, *CRB3*, *JUP*, *LAYN*, *MAPK13, MPZL1*, *MSN*, *SYK*, and *YWHAH*. EMT is associated with the loss of apical to basal cell polarity, the down-regulation of genes encoding proteins maintaining epithelial cell–cell adhesions, such as tight junction proteins, and the up-regulation of genes encoding proteins promoting cell migration and invasion. The DE genes encoding cell-cell adhesion junction molecules within the AJ term were *CLDN7*, *CRB3*, and *JUP*, which were each down regulated in the *BAP1* mutants. In addition, *CGN* and *CAMSAP3*, while not included in the AJ Hallmark gene set, are part of the apical junction complex and were also down-regulated. *MAPK13* (down-regulated) encodes a member of the p38 MAP kinase family. p38 MAPK inhibition has been correlated with resistance to anoikis, which can increase the survival of circulating tumor cells [42]. The up-regulated genes in the AJ hallmark included *LAYN*, *MPZL1* and *MSN*, which promote cell spreading, membrane protrusions, migration, invasion and/or EMT [43,44]. Five other genes of interest to the UM field were differentially expressed (p adj < 0.05) in the same direction in both mouse and human UM. These included *FABP5* [45], *FZD6* [45], *HTR2B* [46,47], *ANO6* [48] and *PROS1* [49] (**Tables S1 and S3).**

Kaplan-Meier plots show that low expression of five of the AJ genes is significantly associated with reduced survival in the TCGA UVM dataset (**Figure 7F**). *JUP* expression was the most significantly associated with outcome (p=6×10^-7^), exceeding that of even *BAP1*. *JUP* encodes the plakoglobin protein. Plakoglobin, a catenin, is a component of both desmosomes and adherens junctions and links cadherins to the actin cytoskeleton. Plakoglobin is closely related to β-catenin, with both proteins containing multiple armadillo motif repeats. However, unlike β-catenin, plakoglobin generally acts as a tumor suppressor in cancer [50].

## DISCUSSION

*BAP1* is a critical tumor suppressor in many different cancer types. *BAP1* loss is strongly associated with metastatic disease and poor clinical outcome in UM. Hence, we anticipated that the loss of *Bap1* would enhance tumor growth and/or dissemination in a mouse model of oncogenic GNAQ-driven UM. Instead, our findings revealed a complex relationship between *Bap1* status and tumor behavior in the model. In the *Mitf-cre*; GNAQ^Q209L^ model, *Bap1^flox/KO^* was associated with a reduction in the number and size of lung lesions. *Bap1* heterozygous loss was associated with melanocyte expansion in structurally abnormal eyes (without GNAQ^Q209L^) and with macroscopic liver lesions (with GNAQ^Q209L^). Finally, transcriptomic analysis identified a shared *BAP1*-associated gene expression program in mouse and human UM that was strongly enriched for epithelial-to-mesenchymal transition (EMT) and apical junction pathways. Together, these findings suggest that Bap1 influences tumor-cell behavior in a manner that depends on gene dosage, tissue context and the surrounding microenvironment.

### No evidence that *Bap1* loss enhances primary melanocytic growth in the GNAQ-driven model

GNAQ^Q209L^ expression in melanocytes was sufficient to produce intraocular melanomagenesis in all mice and all *Bap1* genotypes examined. Therefore, this oncogenic driver is a strong determinant of primary tumor initiation/growth in this model. Our analysis of the role of *Bap1* loss in intraocular tumors was confounded by microphthalmia caused by *Mitf-cre*. However, there was no evidence to support increased primary melanocytic growth in the meninges of the central nervous system in any of the three *Bap1* mutant genotypes [51]. In this mouse model, a large number of melanocytes initiate GNAQ^Q209L^ expression collectively, and they may be able to support each other’s growth or limit the effects of natural tumor suppressing mechanisms that necessitate additional gene mutations in spontaneous/naturally occurring melanoma.

### *Bap1^flox/KO^* unexpectedly reduces lung lesion burden

A striking finding in this study was the reduction in the number of lung lesions in *Bap1^flox/KO^* mice. This genotype is akin to UM arising in the rare human disorder, BAP1 tumor predisposition syndrome, in which individuals inherit a *BAP1* focal mutation. They are at increased risk of developing UM, following the same molecular path as sporadic *BAP1*-mutant UM (*i.e.* GNAQ or GNA11 activation plus monosomy 3) [52]. The *Bap1^flox/KO^*animals carry one constitutive *Bap1*-null allele and one *Bap1 flox* allele. Consequently, the reduced lung lesion burden observed in *Bap1^flox/KO^* animals could reflect a melanocyte-cell-autonomous consequence of complete *Bap1* loss, an effect of constitutive *Bap1* haploinsufficiency in the surrounding environments (eye and/or lung), or an interaction between these two effects. *Bap1^KO^/+* mice did not exhibit this phenotype, so constitutive *Bap1* haploinsufficiency in the surrounding environments alone is not sufficient to produce it. In people with BAP1 tumor predisposition syndrome, the UM metastatic risk is similar to sporadic *BAP1*-mutant UM, according to a recent study [53].

Importantly, the lung lesions in the *Mitf-cre*; GNAQ^Q209L^ model have not been definitively established as metastases. Thus, changes in lung lesion burden cannot be interpreted unequivocally as changes in metastatic dissemination. The lung is not the most common site of metastasis in human UM. Unlike many solid tumors, UM exhibits a striking tropism for the liver, while isolated pulmonary metastases are relatively uncommon. Thus, the abundant lung lesions observed in the *Mitf-cre*; GNAQ^Q209L^ model may not faithfully recapitulate human UM. On the other hand, small lesions like the lung lesions in the mouse model would likely remain clinically undetected in people, as metastases are typically identified only once they reach a size detectable by imaging or become symptomatic. Thus, the apparent discrepancy between the mice and human clinical observations could also reflect differences in detection thresholds and time scales.

### *Bap1* haploinsufficiency is associated with liver lesions

*Bap1* heterozygosity was associated with the appearance of macroscopic liver lesions in the *Mitf-cre*; GNAQ^Q209L^ mouse model. Although the number of animals with liver lesions was small, this finding was reproduced in an independent breeding cohort. The liver lesions are particularly striking because hepatic metastasis is the defining site of dissemination in human UM. Although we cannot conclude that the liver lesions are metastases, their melanocytic phenotype and replacement growth pattern are compatible with hepatic UM metastases. This pattern is characteristically made up of individual S100B-positive tumor cells intermingled with hepatocytes, suggesting that these lesions interact closely with the surrounding liver. The prominent accumulation of melanin-containing CD68-positive macrophages is particularly striking. Whether these macrophages are tumor-promoting, tumor-restraining, or functionally heterogeneous remains to be determined. The observation that mice with liver lesions had larger lung lesions without having more lung lesions raises the possibility that hepatic tumor growth could influence systemic disease progression, although the small number of animals and observational nature of this analysis prevent a causal conclusion.

We also identified an unusual Arg1-positive, highly proliferative cell population within some liver lesions that may represent an immature myeloid population. The morphology and Arg1 expression are compatible with, but do not establish, an MDSC-like identity. Definitive identification will require additional lineage and maturation markers. Nevertheless, the finding raises the possibility that Bap1-associated liver lesions may develop within a distinctive immunological microenvironment and could provide a model for studying interactions between melanoma cells and hepatic myeloid populations.

### *Mitf-cre; Bap1^flox^/+* in the absence of GNAQ^Q209L^ reveals a context-dependent effect on uveal melanocytes

Our observation that *Mitf-cre/+; Bap1^flox^/+* eyes contain expanded pigmented, S100B-positive and RPE65-negative tissue provides another indication that Bap1 function is dependent on tissue context. Importantly, this phenotype was not observed in *Bap1^KO^/+* eyes or in *Tyr-cre/+; Bap1^flox^/+* eyes. Thus, heterozygous *Bap1* loss alone is insufficient to produce the phenotype and instead appears to interact specifically with the consequences of *Mitf-cre* and the associated abnormal ocular architecture. One explanation is that the retinal and RPE abnormalities produced by *Mitf-cre* alter the physical environment in which uveal melanocytes reside. Loss of the retina and Bruch’s membrane in areas of the most extensive melanocyte expansion is consistent with a model in which normal tissue architecture constrains melanocyte growth, while disruption of these barriers creates a permissive environment for expansion or invasion. This is potentially relevant to human UM, in which tumor cells that breach Bruch’s membrane can expand into the subretinal space and develop characteristic mushroom-shaped growth patterns [40].

### *BAP1* loss produces a conserved transcriptional program associated with EMT and cell adhesion

The RNA-seq analysis of primary UM tissue provides a molecular perspective on these phenotypes. We chose to use the *Bap1^flox^*^/^+ mouse genotype, as it was cleanest to interpret and was associated with liver involvement. Both mouse and human *BAP1*-deficient UM exhibited thousands of differentially expressed genes, demonstrating that *BAP1* loss is associated with a broad transcriptional change rather than a small number of isolated downstream effects. Despite the differences between the mouse model and human tumors, 270 genes were differentially expressed in the same direction in both species, significantly more than expected by chance. This conserved subset therefore provides a useful starting point for identifying biological processes associated with *BAP1* loss. The strongest pathway enrichment among these conserved genes was the EMT hallmark, with the majority of shared EMT-associated genes increased in *BAP1*-deficient tumors. ZEB1, a major regulator of EMT, was also increased. The accompanying changes in apical junction genes provide a complementary indication of altered cell-cell adhesion and polarity. In particular, down-regulation of *CLDN7*, *CRB3*, *JUP*, *CGN*, and *CAMSAP3* is consistent with disruption of epithelial-like junctional organization, while increased expression of genes such as *LAYN*, *MSN*, and *MPZL1* is consistent with changes in cell spreading, membrane organization, migration, or invasion. Although UM is not an epithelial cancer and therefore does not undergo classical EMT in the same way as carcinomas, the EMT Hallmark gene set can capture a broader transition toward a more motile, less adhesive, and more plastic cellular state. We therefore interpret the enrichment primarily as evidence for an EMT-like transcriptional program rather than literal epithelial-to-mesenchymal conversion. Such a program could be particularly relevant to the ability of *BAP1*-deficient UM cells to alter their interactions with tissue structures and to migrate or survive in different microenvironments.

The association of several apical junction genes with patient outcome further supports the potential biological importance of this pathway. *JUP*, which encodes plakoglobin, was especially notable. Reduced *JUP* expression was strongly associated with poorer survival in the TCGA UVM cohort, even more strongly than *BAP1* itself in this analysis. Because plakoglobin participates in both desmosomal and adherens junction complexes and links cadherins to the cytoskeleton, its loss provides a reasonable connection between BAP1-associated transcriptional changes and altered cell-cell adhesion [50]. The survival analysis is associative and functional experiments will be required to determine whether JUP or other components of the apical junction pathway contribute directly to the aggressive behavior of *BAP1*-deficient UM.

Polycomb repressive complex 1 (PRC1) is an epigenetic repressive complex that contributes to transcriptional silencing, in part through histone H2A-K119 ubiquitination. Thus, PRC1 activity can help maintain certain cell-state programs in a repressed state. A study by Bakhoum *et al* investigated the role of genetic and epigenetic changes in high-risk versus low-risk UM cell lines [14]. Interestingly, RNAseq experiments found that EMT was up-regulated in the high-risk UM cells compared to the low-risk cells and also was up-regulated when PRC1 was inhibited using PRT4165 in the low-risk UM cells. Eight genes were listed in the supplement to support the EMT term, and four of these genes, *SPARC*, *SERPRINE2*, *TNFRSF12A* and *CALU*, were also up-regulated in our shared *BAP1*-mutant EMT signature. In tumors, cancer cells may adopt an intermediate EMT state that increases their motility while retaining cellular plasticity [54]. Intermediate EMT cells have been shown to be more aggressive in several experimental systems [55,56]. Therefore, it is possible that varying degrees of EMT activation might be related to the non-linear *BAP1* gene dosage effects observed in the *Mitf-cre*; GNAQ^Q209L^ model.

### Evolutionary perspectives on EMT in melanoma

The conservation between mouse and humans with regards to *BAP1* appears to lie at the level of biological potential rather than specific oncogenic outputs. EMT represents a deeply conserved developmental program in neural crest cells [57]. Its enrichment signature across species despite the limited overall overlap in differentially expressed genes suggests convergence at the level of a regulatory state [58]. One possible explanation is that *BAP1* loss perturbs chromatin organization in a way that lowers the barrier to activating EMT-associated regulatory modules. Distinct species-specific transcriptional changes may converge on a shared network architecture defined by core EMT regulators and downstream effectors.

### Implications for the role of BAP1 in UM

Taken together, our results suggest a model in which *BAP1* loss alters the state and environmental responsiveness of uveal melanocytes and UM cells rather than simply increasing proliferation. The consequences of this altered state depend on gene dosage and tissue context. This model provides a possible explanation for why *BAP1* deficiency is such a strong prognostic marker in human UM without necessarily behaving as a universal driver of tumor cell proliferation. *BAP1* loss may increase cellular plasticity and the capacity to interact with, adapt to, or invade particular tissue environments. The outcome of this change would then depend on the organ, the stage of tumor progression, the degree of *BAP1* loss, and the surrounding microenvironment.

Several limitations should be considered. The number of animals with liver lesions was small, and the immune-cell analyses were exploratory. The origin of the lung and liver lesions remains unresolved, which limits the extent to which they can be interpreted as metastases. Differential expression and pathway enrichment identify associations rather than mechanisms. Despite these limitations, the combination of genetic, phenotypic, and transcriptomic analyses reveals an unexpected complexity in the relationship between BAP1 and UM progression. Rather than functioning in a simple linear model where less BAP1 always indicates a more aggressive tumor, we found context-dependent phenotypes. The liver phenotype was particularly notable because hepatic metastasis is the defining site of dissemination in human UM. The conserved changes in EMT-like and apical junction programs provide a potential molecular framework for understanding how *BAP1* deficiency could alter tumor-cell plasticity, tissue invasion, and metastatic behavior.

## ACKNOWLEDGEMENTS

We thank Dr. Susan Marshall for providing the mutant mouse line Bap1<tm1a(EUCOMM)Hmgu>/WtsiIeg, INFRAFRONTIER/EMMA (www.infrafrontier.eu, PMID: 25414328), and Helmholtz Zentrum Muenchen from which the mouse line was distributed (EMMA mouse ID:08891). We thank Dr. Gregory Barsh (Department of Genetics, Stanford University) for the gift of the *Mitf-cre* mice. This work was supported by Life Sciences Institute Cores (Bioinformatics, LSI IMAGING, and the Centre for Disease Modeling), which was supported by the UBC GREx Biological Resilience Initiative. Funding for this research was provided by a grant from the Canadian Institutes of Health Research (CIHR), PJT-178178, to C.D. Van Raamsdonk.

## AUTHOR CONTRIBUTIONS

C.V.R. conceived the idea and designed the present work. R.F., F.J., A.N.L, A.L. and C.V.R. carried out the experiments and analyzed the results. R.F., A.N.L and C.V.R. drafted the paper.

## SUPPLEMENTARY FIGURE LEGENDS

**Figure S1. The ‘knockout-first’ allele design for *Bap1* allows for the production of three different alleles.**

The original “*Bap1^tm1a^*” allele contains an IRES:lacZ trapping cassette, neo cassette, and two loxP sites inserted into *Bap1* intron 5 at position 31256950 of chromosome 14 (build GRCm39). A third loxP site is located downstream of Exon 12 at position 31256950. *Bap1^tm1a^* expresses LacZ and disrupts *Bap1* transcription. Removal of the Frt flanked sequences restores *Bap1* expression and creates a Cre-LoxP conditional allele, called the *Bap1^tm1c^* allele, or “*Bap1^flox^*” in this paper. We crossed *Bap1^tm1a^*/+ mice to *PGK-FLP*/+ mice to generate the *Bap1^flox^* allele. One *Bap1^flox^*/+ mouse was then crossed to a *EIIa-cre*/+ mouse to delete exons 6-12 in the germline and create the *Bap1^tm1d^* allele, or “*Bap1^KO^*” in this paper.

**Figure S2. Homozygosity for the *Bap1^KO^* allele is lethal.**

To check that the *Bap1^KO^* allele behaves like a null allele and is lethal when homozygous, we intercrossed *Bap1^KO^*/+ male and female mice. We obtained no *Bap1^KO/KO^* progeny when genotyping at weaning age (expected frequency = 25%, p = 0.0175, chi square analysis).

**Figure S3. *Bap1* deficiency did not affect primary melanocyte overgrowth in the brain meninges at 20 weeks.**

**A)** Representative brains from 20 week old *Mitf-cre*/+; GNAQ^Q209L^ mice with the indicated *Bap1* genotypes, along with a normal brain on the far right for comparison. The dorsal surfaces are shown in the top row and the ventral surfaces are shown in the bottom row. **B)** Quantification of the brain lesions at 20 weeks of age. The dorsal and ventral lesions were summed together. Each dot represents the measurement from one brain. Error bars represent S.E.M. There were no significant differences between the *Bap1* mutant genotypes. *Mitf-cre*/+ mice not expressing GNAQ^Q209L^ did not exhibit any pigmented brain lesions.

**Figure S4. *Bap1* deficiency did not affect primary melanocyte overgrowth in the spine meninges at 20 weeks.**

**A)** Representative images of 20 week old *Mitf-cre*/+; GNAQ^Q209L^ mice with the trunk skin removed, from the indicated *Bap1* genotypes. Dark lesions are associated with the spine. A normal mouse on the far right is shown for comparison. **B)** Quantification of the lesions associated with the spine at 20 weeks of age. Each dot represents the measurement from one mouse. Error bars represent S.E.M. There were no significant differences between the *Bap1* mutant genotypes. *Mitf-cre*/+ mice not expressing GNAQ^Q209L^ did not exhibit any pigmented lesions associated with the spine.

**Figure S5. Immunohistochemistry of a liver lesion in a *Bap1* heterozygous *Mitf-cre;GNAQ^Q209L^* mouse.**

Serial sections of a liver lesion with surrounding normal liver are shown after IHC for **A)** S100b tumor cells, **B)** CD68 macrophages, **C)** CD19 B cells, **D)** Ly6g neutrophils, and **E)** CD3 T cells. Sections were bleached of melanin and counterstained with hematoxylin. The green boxed area on the left is shown at higher magnification on the right.

**Figure S6. Conditional heterozygous loss of *Bap1* via *Tyr-cre* did not produce a pigmentation phenotype in the eye.**

**A)** 11 week old *Tyr-cre/+*; *Bap1^flox^/+* and *Tyr-cre/+*; +/+ mouse eyes were sectioned and stained with H&E. **B)** Graph of the area of pigmentation in the eyes of *Tyr-cre/+*; *Bap1^flox^/+* and *Tyr-cre/+*; +/+ mice. Each point represents the measurement from one eye. Error bars show standard error of the mean. There is no significant difference between these genotypes.

**Figure S7. There was no significant difference in the levels of *Mitf* 1B containing transcripts in UM from the *Mitf-cre*;GNAQ^Q209L^ versus *Tyr-cre*; GNAQ^Q209L^ mouse models.**

**A)** (top) The mouse *Mitf* gene is shown, not to scale. *Mitf* has multiple alternative start exons. 1A and 1M are used in melanocytes. 1M is melanocyte specific. 1H, 1D and 1B are used by the retina pigment epithelium (RPE) of the eye. Except for exon 1M, which is spliced directly to exon 2, all other upstream exons give rise to transcripts that include exon 1B as the common link to exon 2. (bottom) The *Mitf-cre* transgene structure is shown, not to scale. 1M was replaced by Cre.

**B)** RNA was extracted from either dissected UM tissue (GNAQ^Q209L^ models) or the whole eye (mild *Mitf-cre* control) and reverse transcribed for qRT-PCR with a primer set that has one primer in Exon1B and one primer in Exon 2. Expression was normalized to the *Hprt1* housekeeping gene. Each point represents one sample, done in triplicate. *Mitf* 1B containing transcripts were not elevated in *Mitf-cre*; GNAQ^Q209L^ UM compared to *Tyr-cre;* GNAQ^Q209L^ UM. The 1B expression observed in UM is from the 1A start exon in the endogenous *Mitf* gene.

**Figure S8. Abnormal splicing products detected in *Bap1* mutant UM RNAseq experiment, related to Figure 6**.

A sashimi plot of *Bap1* transcripts from the RNA sequencing of *Mitf-cre*/+; *R26-fs-GNAQ^Q209L^*/+ uveal melanoma that was either *Bap1* +/+ (WT) or *Bap1^flox^*/+ (Mut). Each sample name is indicated on the right hand side. Samples 3-WT and 4-WT were originally sequenced as two technical replicates each (1,2) and were subsequently collapsed for DE analysis. *Bap1* exons 4-17 are shown in the map at the bottom. Arrows point out reads that splice the end of exon 5 to exon 13. The black number associated with the arrow is the number of times this read was represented in the sequencing run. The *Bap1* mutant allele deletes exons 6-12. No other abnormal splices events were found.

## SUPPLEMENTARY TABLE LEGENDS

**Table S1.** Results of DESeq2 analysis of differentially expressed genes in *Bap1* mutant versus wildtype mouse UM.

**Table S2.** Features of the TCGA-UVM cases used for analysis.

**Table S3.** Results of DESeq2 analysis of differentially expressed genes in *BAP1* mutant versus wildtype human UM.

**Table S4.** Lists of genes that are either up-regulated, down-regulated, or not DE in *Bap1*-deficient mouse UM and *BAP1*-deficient human UM.

**Table S5.** Pathway analysis results of the 270 shared DE gene set in *Bap1* mutant mouse and *BAP1* mutant human UM.

**Table S6.** Primers used to distinguish the *Bap1* alleles used in this study.

