## Supplementary Figures S1-S8 for "BAP1 deficient human and mouse uveal melanomas up-regulate a shared EMT pathway"

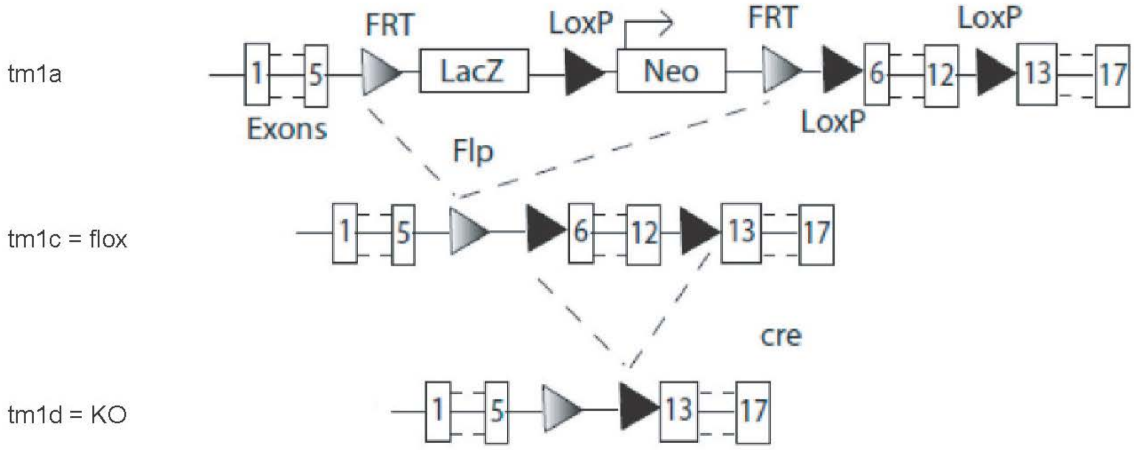


**Figure S1. The 'knockout-first' allele design for *Bap1* allows for the production of three different alleles.**

The original "*Bap1*$^{tm1a}$" allele contains an IRES:lacZ trapping cassette, neo cassette, and two loxP sites inserted into *Bap1* intron 5 at position 31256950 of chromosome 14 (build GRCm39). A third loxP site is located downstream of Exon 12 at position 31256950. *Bap1*$^{tm1a}$ expresses LacZ and disrupts *Bap1* transcription. Removal of the Frt flanked sequences restores *Bap1* expression and creates a Cre-LoxP conditional allele, called the *Bap1*$^{tm1c}$ allele, or "*Bap1*$^{flox}$" in this paper. We crossed *Bap1*$^{tm1a}$/+ mice to *PGK-FLP*/+ mice to generate the *Bap1*$^{flox}$ allele. One *Bap1*$^{flox}$/+ mouse was then crossed to a *EIIa-cre*/+ mouse to delete exons 6-12 in the germline and create the *Bap1*$^{tm1d}$ allele, or "*Bap1*$^{KO}$" in this paper.

| *Allele* | *Bap1$^{KO/KO}$* (25% expected) | *Bap1$^{KO/+}$* (50% expected) | *Bap1$^{+/+}$* (25% expected) |
|---|---|---|---|
| Male | 0 | 7 | 6 |
| Female | 0 | 6 | 3 |
| Total | 0 (0%) | 13 (59.09%) | 9 (40.9%) |

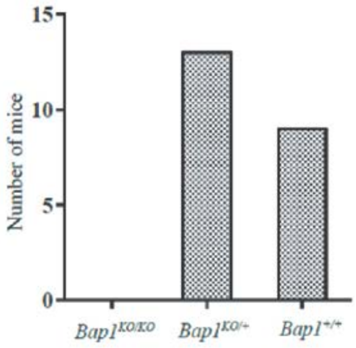


**Figure S2. Homozygosity for the *Bap1$^{KO}$* allele is lethal.**

To check that the *Bap1$^{KO}$* allele behaves like a null allele and is lethal when homozygous, we intercrossed *Bap1$^{KO}$*/+ male and female mice. We obtained no *Bap1$^{KO/KO}$* progeny when genotyping at weaning age (expected frequency = 25%, $p = 0.0175$, chi square analysis).

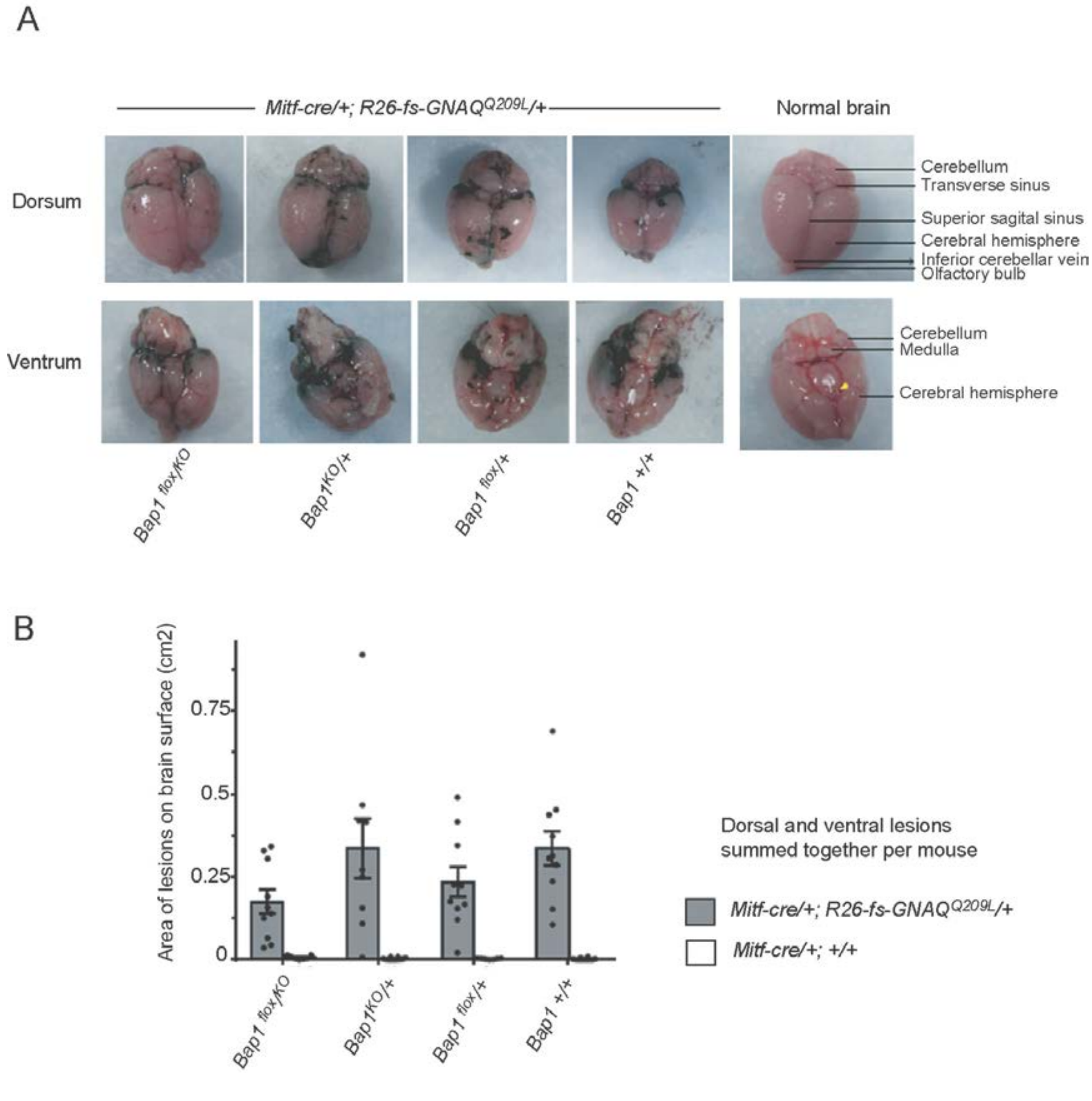


**Figure S3. *Bap1* deficiency did not affect primary melanocyte overgrowth in the brain meninges at 20 weeks.**

**A)** Representative brains from 20 week old *Mitf-cre*/+; GNAQ$^{Q209L}$ mice with the indicated *Bap1* genotypes, along with a normal brain on the far right for comparison. The dorsal surfaces are shown in the top row and the ventral surfaces are shown in the bottom row. **B)** Quantification of the brain lesions at 20 weeks of age. The dorsal and ventral lesions were summed together. Each dot represents the measurement from one brain. Error bars represent S.E.M. There were no significant differences between the *Bap1* mutant genotypes. *Mitf-cre*/+ mice not expressing GNAQ$^{Q209L}$ did not exhibit any pigmented brain lesions.

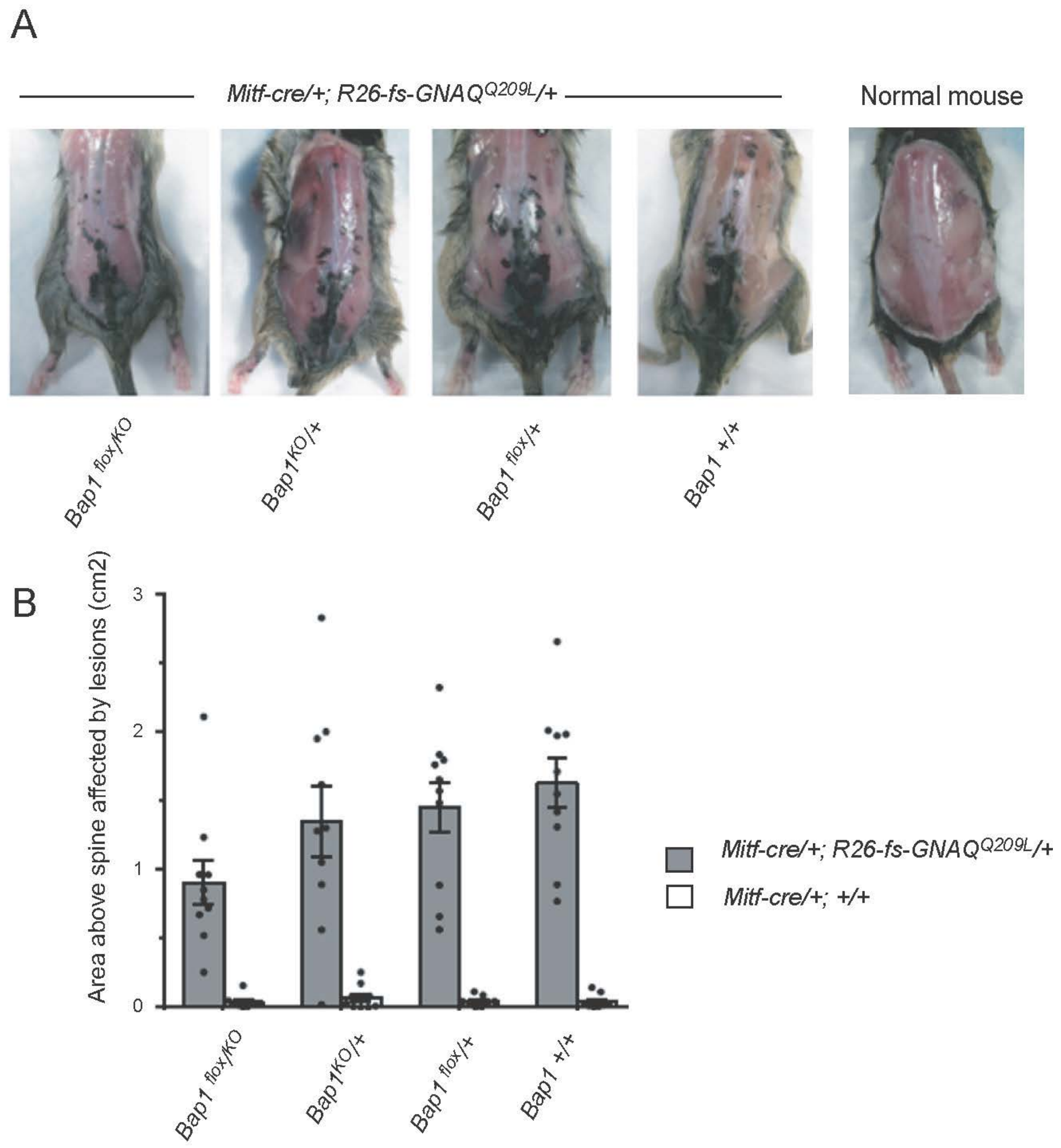


**Figure S4. *Bap1* deficiency did not affect primary melanocyte overgrowth in the spine meninges at 20 weeks.**

**A)** Representative images of 20 week old *Mitf-cre*/+; GNAQ$^{Q209L}$ mice with the trunk skin removed, from the indicated *Bap1* genotypes. Dark lesions are associated with the spine. A normal mouse on the far right is shown for comparison. **B)** Quantification of the lesions associated with the spine at 20 weeks of age. Each dot represents the measurement from one mouse. Error bars represent S.E.M. There were no significant differences between the *Bap1* mutant genotypes. *Mitf-cre*/+ mice not expressing GNAQ$^{Q209L}$ did not exhibit any pigmented lesions associated with the spine.

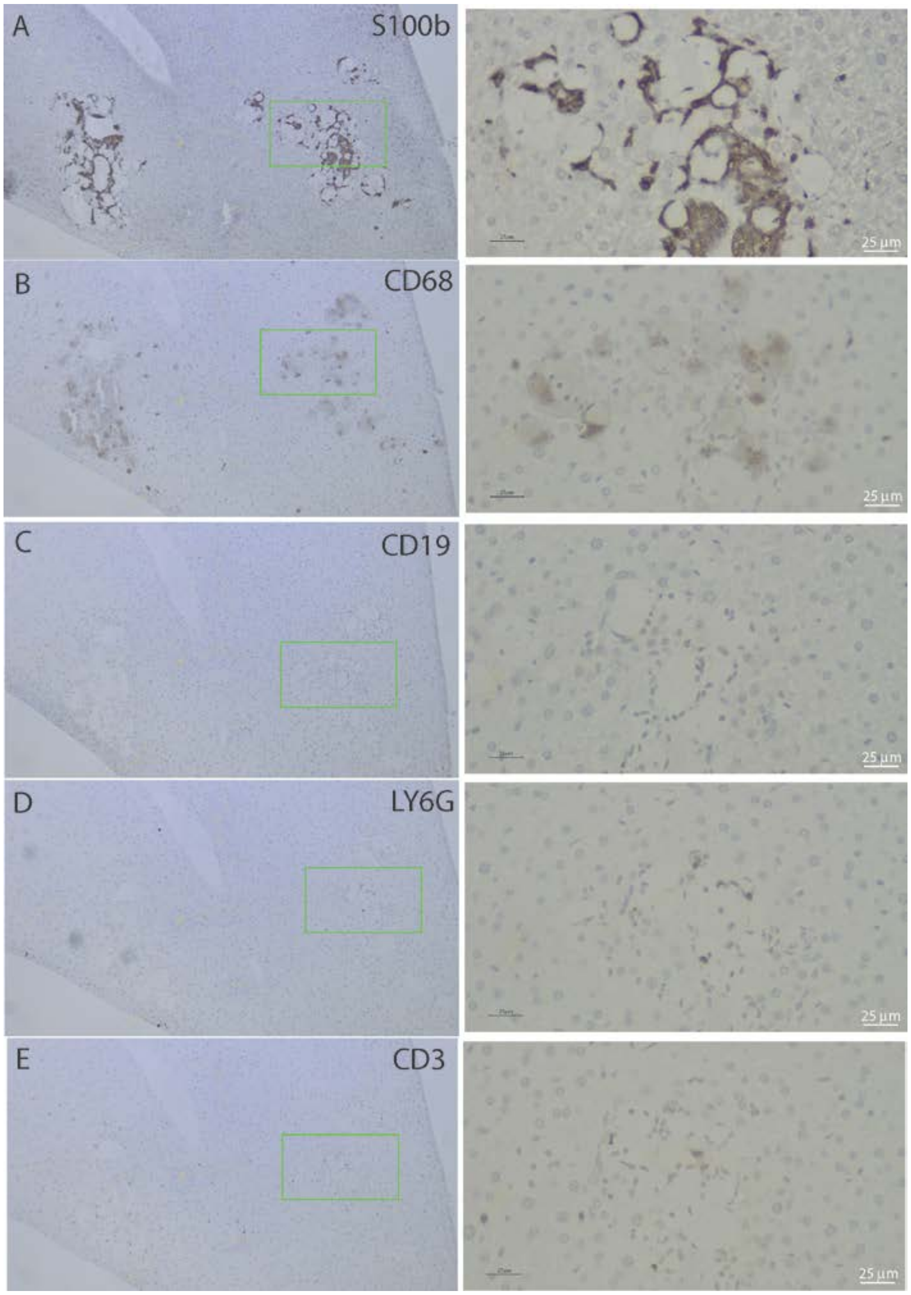


**Figure S5. Immunohistochemistry of a liver lesion in a *Bap1* heterozygous *Mitf-cre;GNAQ*$^{Q209L}$ mouse.**

Serial sections of a liver lesion with surrounding normal liver are shown after IHC for **A)** S100b tumor cells, **B)** CD68 macrophages, **C)** CD19 B cells, **D)** Ly6g neutrophils, and **E)** CD3 T cells. Sections were bleached of melanin and counterstained with hematoxylin. The green boxed area on the left is shown at higher magnification on the right.

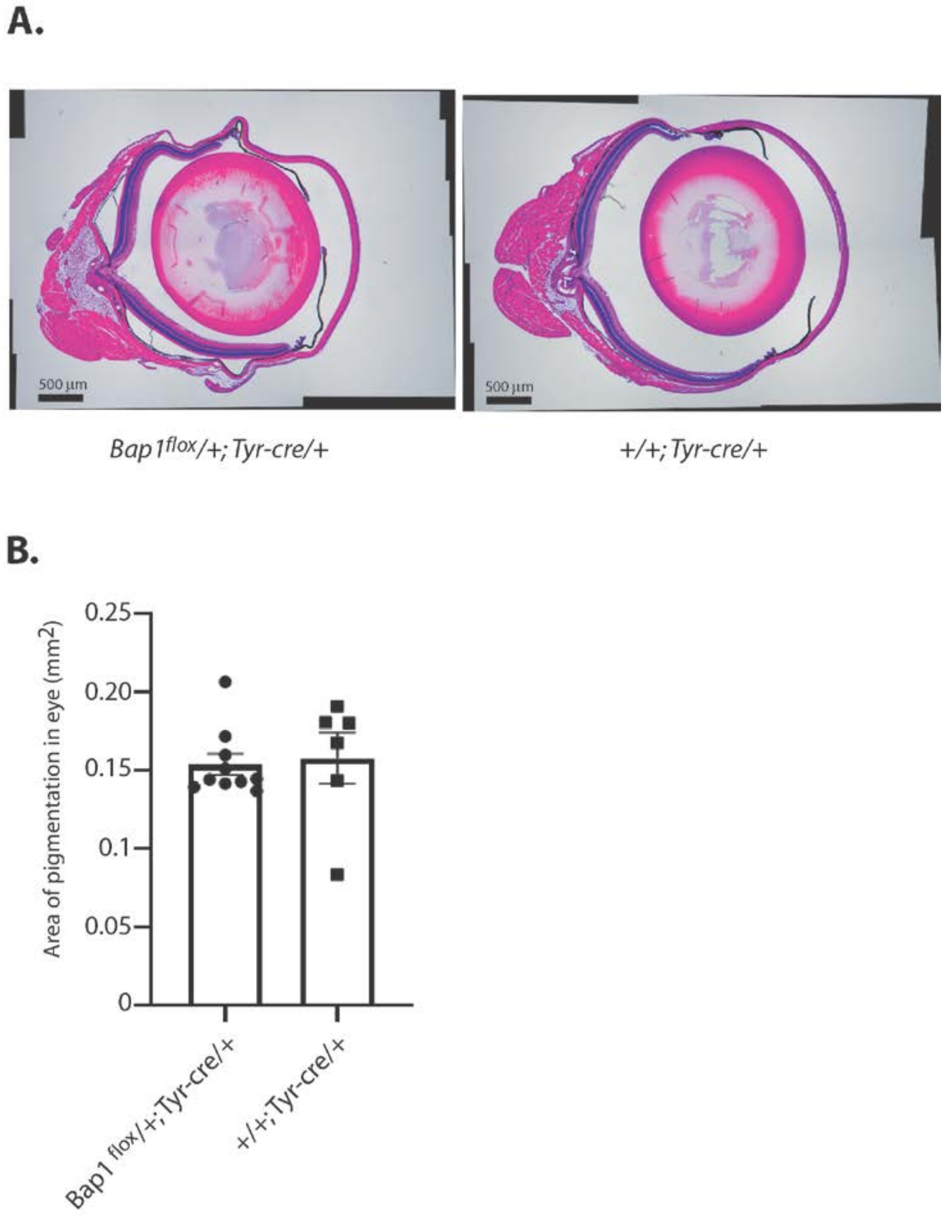


**Figure S6. Conditional heterozygous loss of *Bap1* via *Tyr-cre* did not produce a pigmentation phenotype in the eye.**

**A)** 11 week old *Tyr-cre*/+; *Bap1*$^{flox}$/+ and *Tyr-cre*/+; +/+ mouse eyes were sectioned and stained with H&E. **B)** Graph of the area of pigmentation in the eyes of *Tyr-cre*/+; *Bap1*$^{flox}$/+ and *Tyr-cre*/+; +/+ mice. Each point represents the measurement from one eye. Error bars show standard error of the mean. There is no significant difference between these genotypes.

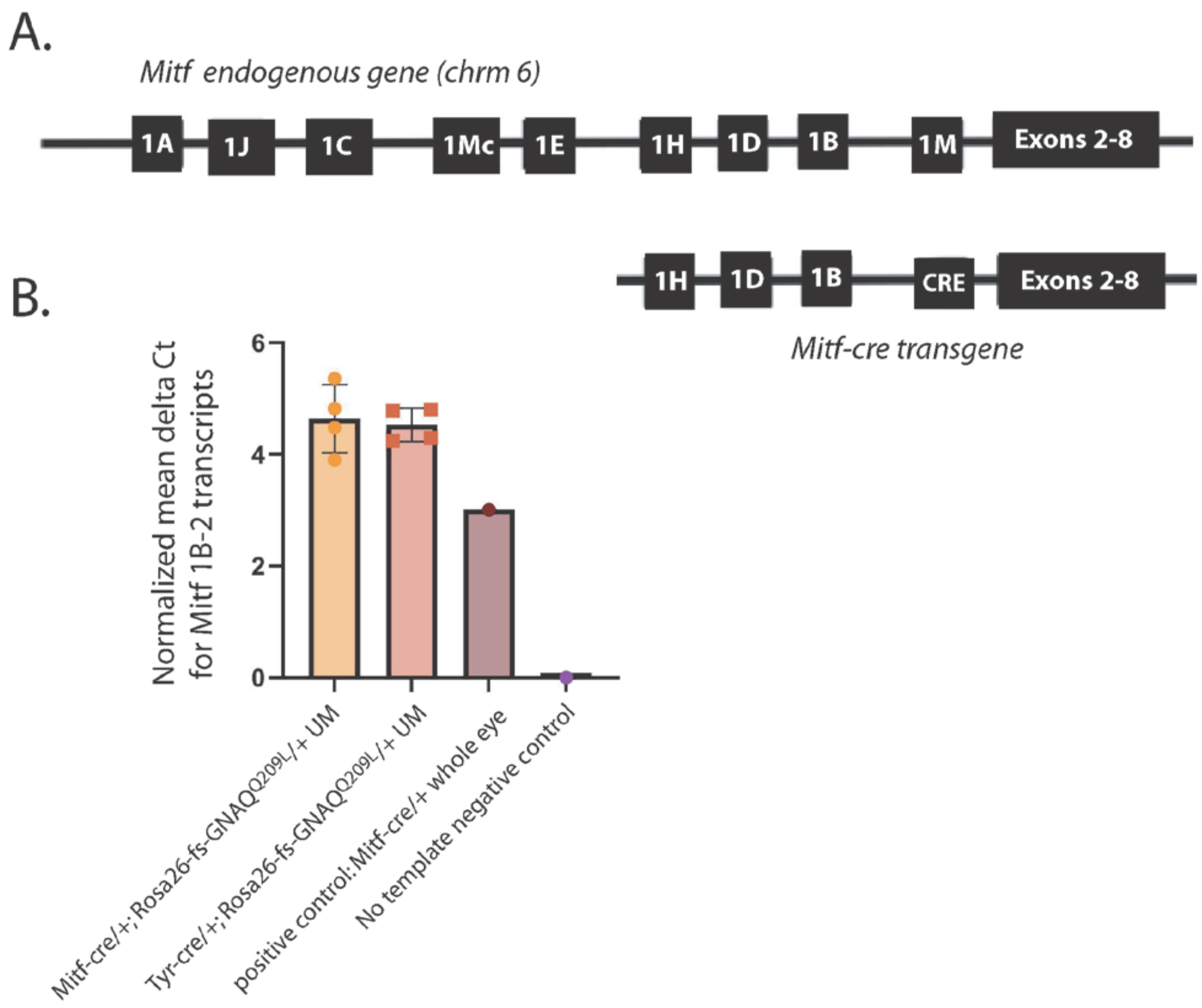


**Figure S7. There was no significant difference in the levels of *Mitf* 1B containing transcripts in UM from the *Mitf-cre*;GNAQ$^{Q209L}$ versus *Tyr-cre*; GNAQ$^{Q209L}$ mouse models.**

**A)** (top) The mouse *Mitf* gene is shown, not to scale. *Mitf* has multiple alternative start exons. 1A and 1M are used in melanocytes. 1M is melanocyte specific. 1H, 1D and 1B are used by the retina pigment epithelium (RPE) of the eye. Except for exon 1M, which is spliced directly to exon 2, all other upstream exons give rise to transcripts that include exon 1B as the common link to exon 2. (bottom) The *Mitf-cre* transgene structure is shown, not to scale. 1M was replaced by Cre.

**B)** RNA was extracted from either dissected UM tissue (GNAQ$^{Q209L}$ models) or the whole eye (mild *Mitf-cre* control) and reverse transcribed for qRT-PCR with a primer set that has one primer in Exon1B and one primer in Exon 2. Expression was normalized to the *Hprt1* housekeeping gene. Each point represents one sample, done in triplicate. *Mitf* 1B containing transcripts were not elevated in *Mitf-cre*; GNAQ$^{Q209L}$ UM compared to *Tyr-cre;* GNAQ$^{Q209L}$ UM. The 1B expression observed in UM is from the 1A start exon in the endogenous *Mitf* gene.

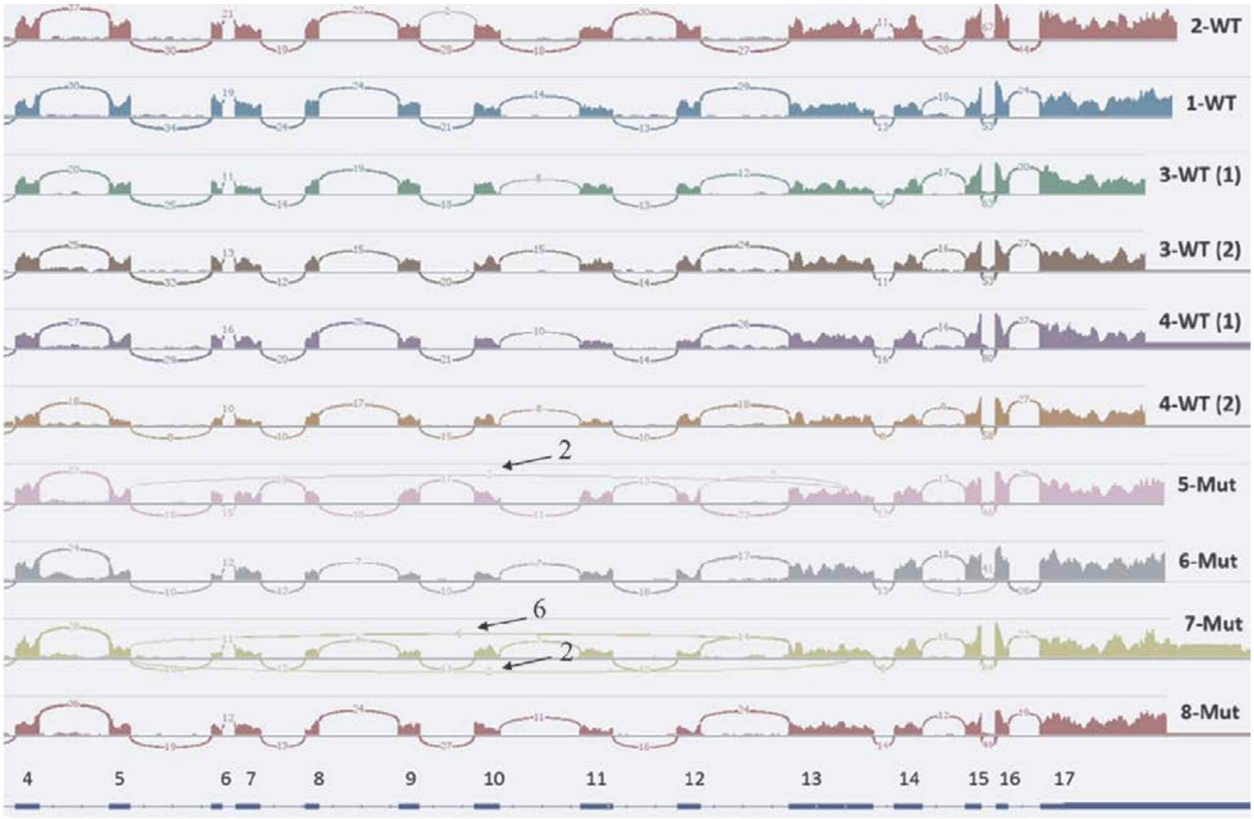


**Figure S8. Abnormal splicing products detected in *Bap1* mutant UM RNAseq experiment, related to Figure 6.**

A sashimi plot of *Bap1* transcripts from the RNA sequencing of *Mitf-cre*/+; *R26-fs-GNAQ*$^{Q209L}$/+ uveal melanoma that was either *Bap1* +/+ (WT) or *Bap1*$^{flox}$/+ (Mut). Each sample name is indicated on the right hand side. Samples 3-WT and 4-WT were originally sequenced as two technical replicates each (1,2) and were subsequently collapsed for DE analysis. *Bap1* exons 4-17 are shown in the map at the bottom. Arrows point out reads that splice the end of exon 5 to exon 13. The black number associated with the arrow is the number of times this read was represented in the sequencing run. The *Bap1* mutant allele deletes exons 6-12. No other abnormal splices events were found.
